# The big tau splice isoform resists Alzheimer’s-related pathological changes

**DOI:** 10.1101/2024.07.30.605685

**Authors:** Dah-eun Chloe Chung, Xue Deng, Hari K. Yalamanchili, Jean-Pierre Revelli, Alexander L. Han, Bakhos Tadros, Ronald Richman, Michelle Dias, Fatemeh Alavi Naini, Steven Boeynaems, Bradley T. Hyman, Huda Y. Zoghbi

## Abstract

In Alzheimer’s disease (AD), the microtubule-binding protein tau becomes abnormally hyperphosphorylated and aggregated in selective brain regions such as the cortex and hippocampus^1–3^. However, other brain regions like the cerebellum and brain stem remain largely intact despite the universal expression of tau throughout the brain. Here, we found that an understudied splice isoform of tau termed “big tau” is significantly more abundant in the brain regions less vulnerable to tau pathology compared to tau pathology-vulnerable regions. We used various cellular and animal models to demonstrate that big tau possesses multiple properties that can resist AD-related pathological changes. Importantly, human AD patients show a higher expression level of pathology-resisting big tau in the cerebellum, the brain region spared from tau pathology. Our study examines the unique properties of big tau, expanding our current understanding of tau pathophysiology. Altogether, our data suggest that alternative splicing to favor big tau is a viable strategy to modulate tau pathology.

## MAIN TEXT

Hyperphosphorylation and abnormal aggregation of the microtubule-binding protein tau are major pathological hallmarks of Alzheimer’s disease (AD) and other tauopathies^2,3^. Tau pathology is strongly and positively correlated with cognitive impairment in AD patients^4^, which highlights its importance to the pathophysiology of AD and emphasizes the need to develop effective tau-targeting therapies. During AD progression, tau pathology spreads throughout the brain and heavily affects selective brain regions like the cortex and hippocampus^1^. Intriguingly, the cerebellum rarely develops tau pathology, even at the most advanced stage of AD^1^ or tauopathies resulting from *MAPT* mutations that promote tau aggregation^5–7^. Similarly, the brainstem is largely spared from tau pathology, except for a few nuclei that bear tau inclusions^8^. The reasons for this regional sparing from tau pathology are currently unknown but hint at the possibility that tau is either cleared more efficiently or resists misfolding and aggregation in these brain regions. While prior studies have examined the levels of tau and phosphorylation-regulating enzymes in these resistant brain regions^9,10^, the question of how they are protected from developing tau pathology remains poorly understood.

Alternative splicing that includes or excludes exons 2, 3, and 10 of the *MAPT* gene leads to six major isoforms of tau^2,3^. These tau isoforms vary in their protein structures depending on whether they include zero, one, or two N-terminal inserts (0N, 1N, or 2N) and either three or four repeats (3R or 4R) in the microtubule-binding regions. Considering these structural differences, tau isoforms are thought to possess varying biochemical properties. For instance, the four-repeat 4R tau isoforms are reported to be more aggregation-prone than the three-repeat 3R tau isoforms^11,12^. Given that many pathogenic tau mutations associated with frontotemporal dementia promote tau exon 10 inclusion and result in a higher ratio of 4R tau^13^, most studies analyzing tau isoforms have focused on 3R versus 4R tau isoforms. Yet, in addition to these six major isoforms, there is another much less-studied tau isoform termed “big tau”, which is generated by the inclusion of *MAPT* exons 4a and 6^14^. As the name “big tau” suggests, this atypical isoform is *much* bigger (758 amino acids) than any of the six major tau isoforms (ranging from 352 to 441 amino acids) due to its substantially elongated N-terminal projection region (NTR). Since its initial identification in the 1990s, big tau has been thought to be the predominant tau isoform in the peripheral nervous system (PNS), hence bearing an alternative name “PNS-tau”^15–18^. So far, big tau has been largely understudied in the central nervous system (CNS) compared to regular tau isoforms, and its biochemical properties or contributions to the pathogenesis of neurodegenerative diseases have remained unexplored.

In this study, we found that the expression level of big tau is significantly elevated in both the cerebellum and brainstem, brain regions spared from overt tau pathology in AD. Importantly, we discovered that big tau does not display the typical AD-related hyperphosphorylation and is more efficiently ubiquitinated and subsequently rapidly degraded. Using cellular and mouse models, we further demonstrated that big tau possesses a significantly enhanced microtubule-binding capacity and diminished aggregation propensity. Lastly, we show that, compared to cognitively normal control individuals, AD patients display a higher level of big tau in the pathology-resistant cerebellum but not in the pathology-sensitive cortex. Thus, we uncover an exciting new aspect of tau pathophysiology by revealing the regional abundance and unique features of big tau. This insight has implications for therapeutic intervention through modulation of tau splicing to favor the big tau isoform.

## Results

### Big tau is highly expressed in the cerebellum and brainstem

Given the perplexing regional brain vulnerability to tau pathology in AD, we first sought to examine the expression pattern of tau isoforms across different brain regions. We performed western blots using protein lysates from the cortex, hippocampus, cerebellum, and brainstem of 4-6-months-old (mo) C57/B6 mice and detected endogenous mouse tau using a total tau antibody (i.e., tau-5). In addition to regular tau isoforms (∼50kDa), we observed tau bands representing a much bigger protein (∼90kDa) expressed in the cerebellum and the brainstem, but not in the cortex and the hippocampus (**Fig. 1a-c**). We also confirmed these ∼90kDa tau bands in the cerebellum and brainstem by using a different total tau antibody (i.e., Dako-tau) and brain lysates extracted from a different mouse strain (i.e., FVB mice) (**Extended Data** Fig. 1a-h). Based on the protein size of tau isoforms described in prior literature^15–18^, we hypothesized the tau bands at a higher molecular weight could represent the “big tau” isoform (**Fig. 1d**), mostly studied in the context of the PNS. To test this hypothesis, we designed primers that can detect the exon junction unique to big tau and measured tau mRNA levels across brain regions. Indeed, quantitative PCR (qPCR) analysis indicated that the big tau transcript was significantly higher in the cerebellum and brainstem compared to the cortex and the hippocampus, while total tau mRNA levels were comparable across these brain regions (**Fig. 1e,f**). Because there was no commercially available antibody specific to the big tau isoform, we generated an antibody that specifically detects the mouse big tau protein (**Extended Data** Fig. 2a-c). With this novel reagent, we confirmed that big tau protein is robustly expressed in the cerebellum and brainstem but minimally in the cortex or hippocampus in mice, coinciding with the original high molecular weight band detected using total tau antibodies (**Fig. 1g,h, Extended Data** Fig. 1i,j). Taken together, our data indicate that big tau is significantly more abundant in both the cerebellum and brainstem.

**Figure 1.**
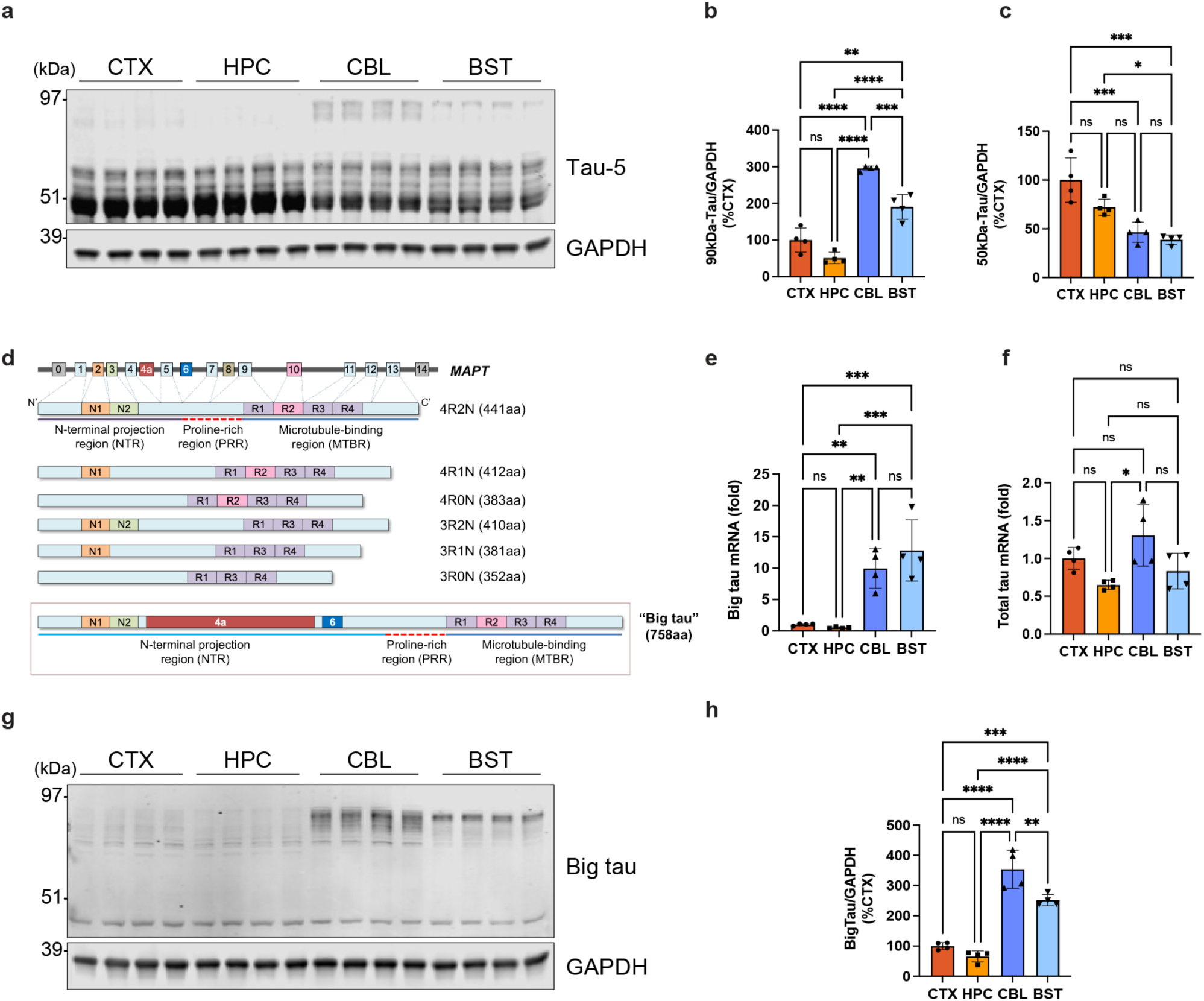
The big tau expression level is significantly elevated in the mouse cerebellum and brainstem. **a-c,** Representative western blots probed with the tau-5 antibody showing tau protein expression across different mouse brain regions. Additional tau bands (∼90kDa) are uniquely present in the cerebellum and brainstem (**a**). Graphs showing the quantification of tau bands at ∼90kDa (**b**) and ∼50kDa (**c**) normalized to GAPDH. **d**, A diagram depicting six regular tau isoforms in the central nervous system (CNS) and big tau (drawn not to scale). Human tau, encoded by the *MAPT* gene, is composed of the N-terminal projection region (NTR) and the C-terminal microtubule-binding region (MTBR), flanked by the proline-rich region (PRR). Alternative splicing of exon 2 (E2), E3, and E10 generates six tau isoforms that range from 352 to 441 amino acids in length. A 758-amino-acid-long tau isoform, termed “big tau” due to its larger protein size, is generated upon the inclusion of E4a and E6 of *MAPT* via alternative splicing. **e-f**, Quantitative PCR (qPCR) results showing the levels of big tau mRNA (**e**) and total tau mRNA (**f**) across different brain regions. **g-h,** Representative western blots showing the expression of big tau protein in the cerebellum and brainstem using a novel mouse big tau antibody (**g**) and their quantification normalized to GAPDH (**h**). All mice were 4-6-month-old (mo) C57/B6. Data (n=4/group) are shown as mean ± SEM. CTX = cortex; HPC = hippocampus; CBL = cerebellum; BST = brainstem; ns = not significant; \**p*<0.05; \*\**p*<0.01; \*\*\**p*<0.001; \*\*\*\**p*<0.0001; one-way ANOVA.

### Big tau is less hyperphosphorylated but more ubiquitinated and degraded

Throughout AD progression, tau undergoes extensive and stereotypic post-translational modifications (PTMs). Specifically, tau is a completely disordered protein with abundant serine and threonine residues that are subject to hyperphosphorylation, particularly during AD pathogenesis. Importantly, this excessive phosphorylation alters the binding of tau to microtubules, a step considered as a key factor promoting its aggregation^19^. Given big tau’s unique structure with an extremely elongated N-terminal projection region (NTR) which is roughly three times longer than that of regular tau (**Fig. 1d**), we hypothesized that big tau could possess biochemical properties distinct from those of regular tau isoforms and that such properties might affect its pathophysiology, including PTM events. In support of our hypothesis, when we performed western blots using brain tissue extracts from young (1-3-mo) vs. old (>12-mo) mice and a phospho-tau antibody (i.e., AT8), we observed that regular isoforms of endogenous mouse tau become robustly hyperphosphorylated in an age-dependent manner (**Fig. 2a-c**). In contrast, we did not detect such phospho-tau signal for big tau in both the cerebellum and brainstem of aged mice (**Fig. 2a-c**), suggesting that big tau is less subject to age-associated hyperphosphorylation.

**Figure 2.**
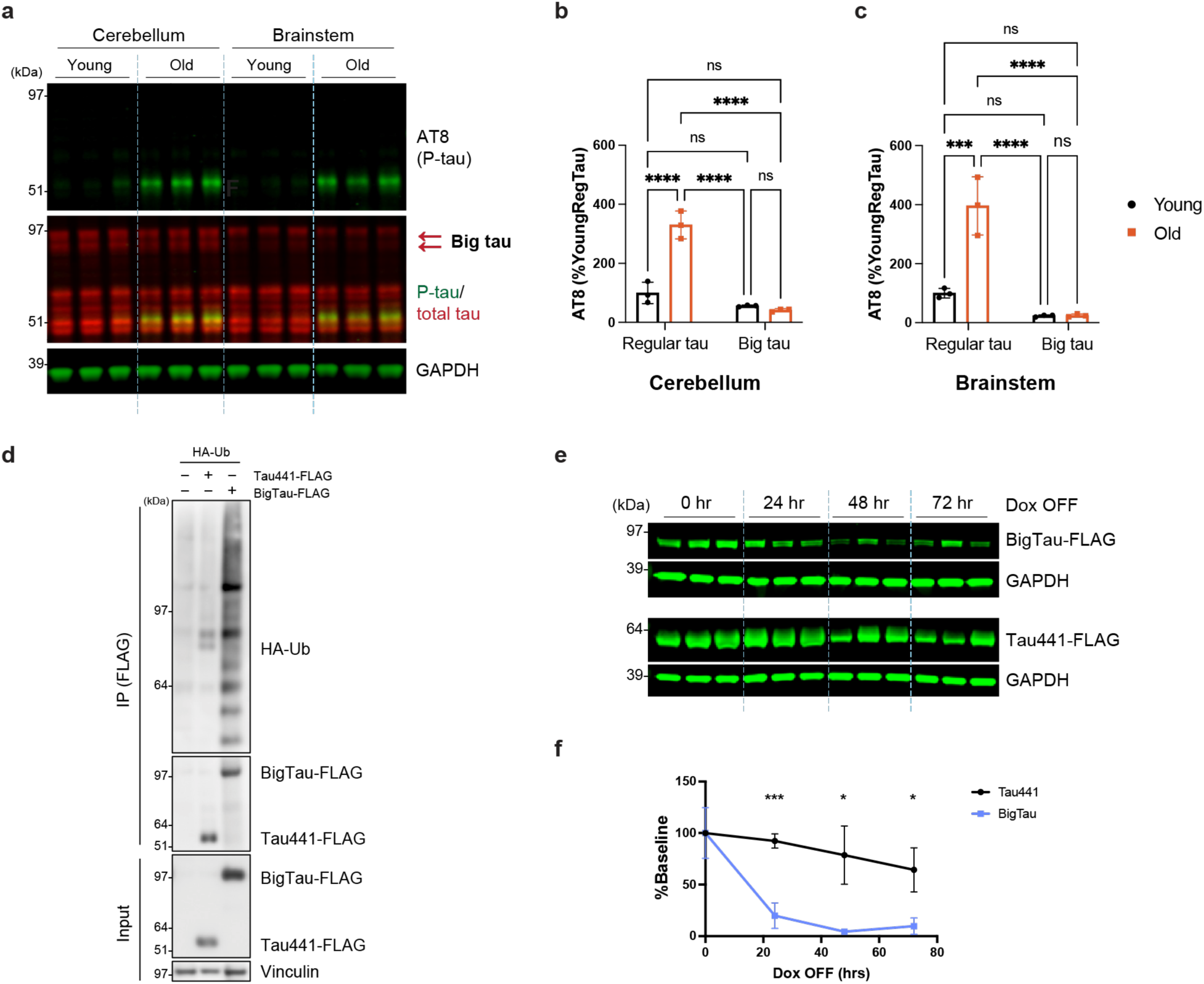
Big tau is less prone to hyperphosphorylation and more susceptible to ubiquitination and degradation. **a-c,** Representative western blots probed with the AT8 antibody showing age-dependent hyperphosphorylation of endogenous mouse tau (**a**) and their quantification for cerebellum (**b**) and brainstem (**c**), both normalized to GAPDH. **d,** Representative western blots probed with the HA antibody showing ubiquitination of FLAG-tagged big tau and tau441 immunoprecipitated (IP) using the FLAG antibody. Vinculin was used as a loading control. **e-f,** Representative western blots probed with the FLAG antibody showing the remaining big tau or tau441 proteins in doxycycline (dox)-inducible stable HEK293T cell lines (**e**). A graph showing the quantification of big tau or tau441 normalized to GAPDH (**f**). Data (n=3/group) are shown as mean ± SEM. P-tau = phosphorylated tau; ns = not significant; \**p*<0.05; \*\*\**p*<0.001; \*\*\*\**p*<0.0001; one-way ANOVA for the mouse data; multiple unpaired t-test for the cell line data.

As is typical for protein aggregates in disease, the hyperphosphorylated tau tangles that accumulate in AD brains are often ubiquitin-positive as the cells try to degrade these pathological assemblies^20^. Tau441 (the longest regular tau isoform; 2N4R [**Fig. 1d**]) has a total of 44 lysine residues that are mostly located in the proline-rich domain and the microtubule-binding domain. In comparison, big tau has 20 additional lysine residues due to its much longer NTR (**Extended Data** Fig. 3). Since lysine residues are the sites of ubiquitination, we examined the degree of ubiquitination of both big tau and tau441 using a ubiquitination assay in HEK293T cells that express HA-tagged ubiquitin and either FLAG-tagged big tau or tau411. By immunoprecipitating (IP) tau isoforms using the FLAG antibody followed by western blots using the HA antibody, we found that big tau was substantially more ubiquitinated than tau441 (**Fig. 2d**). Given that ubiquitination is a prerequisite for protein degradation, we directly compared the protein degradation rates of big tau and tau441 using stable HEK293T cell lines that express doxycycline (Dox)-inducible, FLAG-tagged big tau or tau441. After inducing transgenic expression with Dox for 24 hrs, we removed Dox from the culture media and analyzed the amount of remaining tau proteins by western blots. Here, we found that big tau has a significantly higher protein degradation rate than tau441 (**Fig. 2e,f**). We additionally confirm this by transiently expressing tau isoforms in HEK293T cells and monitoring their levels over time via western blot analyses after treatment with cycloheximide to inhibit the translation of new proteins. Similar to the results in Dox-inducible cell lines, we observed that big tau degrades much faster than tau441 in this cycloheximide chase assay (**Extended Data** Fig. 4). These results, consistent with our finding on big tau’s enhanced ubiquitination (**Fig. 2d**), suggest that the turnover rate of big tau is significantly more rapid. Altogether, our data demonstrate that big tau has a decreased susceptibility to AD-related pathological changes, including hyperphosphorylation and the accumulation of non-degraded ubiquitinated species.

### Big tau has an enhanced microtubule-binding capacity

Since disturbances in the canonical function of tau binding to microtubules have been implicated in its aggregation propensity, we sought to evaluate the microtubule-binding capacity of both big tau and tau441. To do so, we performed ultracentrifugation-mediated microtubule fractionation using Taxol-stabilized lysates of HEK293T cells that express either FLAG-tagged big tau or tau441. After isolating the supernatant that represents the “unbound” fraction and the pellet that represents the “microtubule-bound” fraction (**Fig. 3a**), we analyzed them to determine the ratio of big tau or tau441 between the two fractions by western blots. Intriguingly, we found that big tau is significantly more enriched in the microtubule fraction compared to tau441 (**Fig. 3b,c**), suggesting that big tau has an enhanced ability to bind to microtubules. Intrigued by this finding, we set out to determine if big tau includes any additional microtubule-binding motifs beyond the well-characterized ones in the C-terminal microtubule-binding region (MTBR)^21,22^ and proline-rich region (PRR)^23^. Specifically, we focused on the NTR of big tau since it is the only region distinct from regular tau isoforms. The initial bioinformatics analysis using the MAPanalyzer tool^24^ predicted several potential microtubule-binding motifs within big tau’s NTR (**Extended Data** Fig. 5), suggesting that big tau may possess additional microtubule-binding motifs compared to regular tau. To determine the presence of microtubule-binding motifs specific to big tau, we performed peptide array overlay assays. In brief, 15-mer peptides tiling the entire big tau sequence were spotted on a nitrocellulose membrane. These membranes were incubated with Taxol-stabilized recombinant microtubules and subsequently probed with a tubulin antibody. In doing so, we detected strong microtubule-binding signals in regions specific to big tau indicative of robust microtubule-binding activities (**Fig. 3d,e**). The degree of these additional microtubule-binding activities within big tau’s NTR was similar to—or even stronger—than that of microtubule-binding motifs in the MTBR and PRR domains, suggesting that big tau possesses additional *bona fide* microtubule-binding motifs that could critically contribute to its increased microtubule binding that we observed in cells.

**Figure 3.**
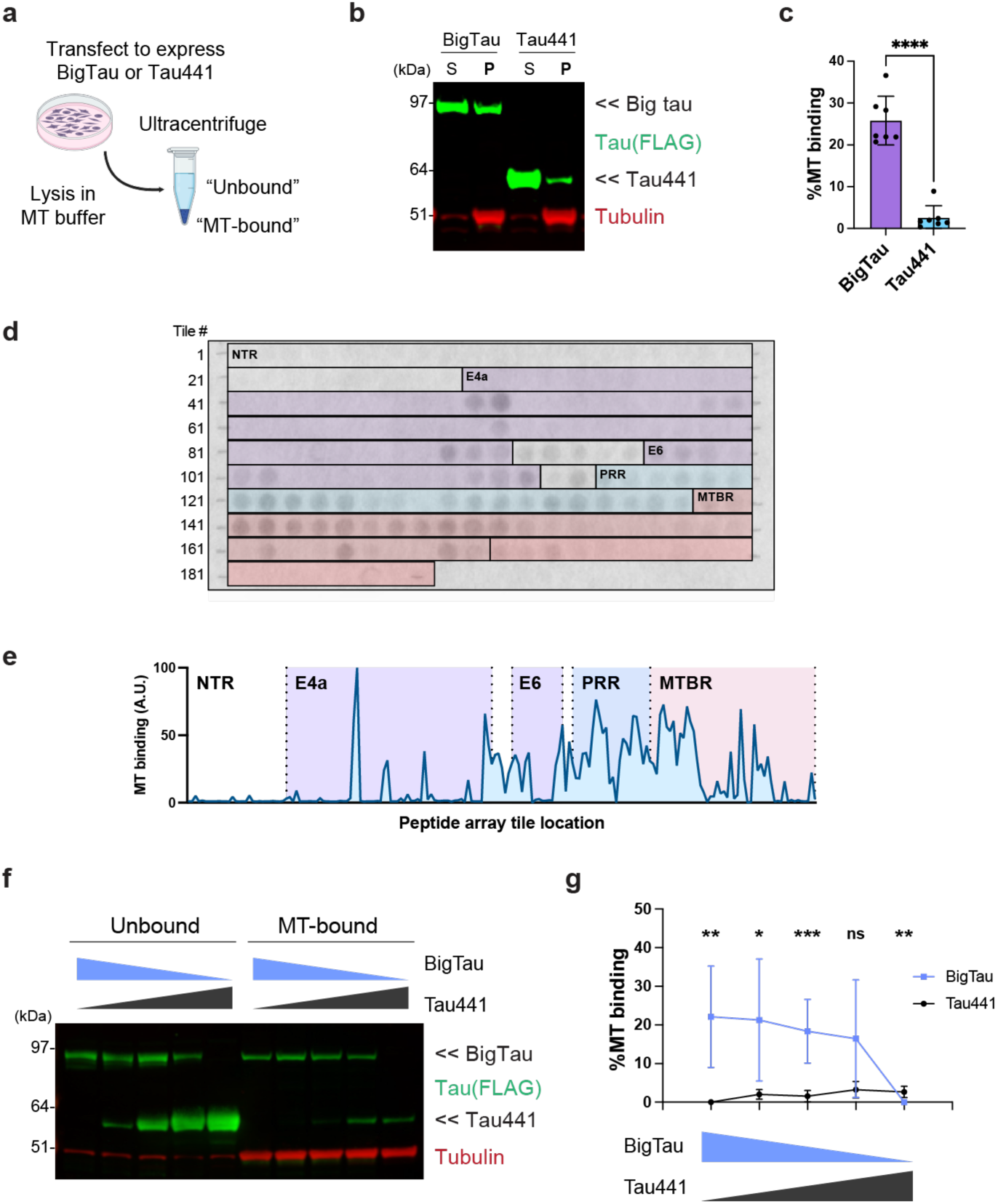
Big tau has a substantially enhanced microtubule-binding capacity via additional microtubule-binding epitopes. **a,** A schematic diagram depicting the ultracentrifuge-mediated microtubule (MT) fractionation of HEK293T cells expressing big tau or tau441 (both tagged with FLAG). **b-c,** Representative western blots probed with the FLAG antibody (**b**) and their quantification (**c**) showing the MT-binding capacity of big tau and tau441. S = supernatant (“unbound” fraction); P = pellet (“MT-bound fraction”). Tubulin antibody was used to ensure the enrichment of MTs in the MT-bound fraction. **d-e,** Peptide array results (**d**) and their quantification (**e**) revealing additional MT-binding activity in regions specific to big tau (generated by exons 4a and 6 of the *MAPT* gene). **f-g,** Representative western blots probed by the FLAG antibody (**f**) and their quantification (**g**) showing big tau outcompeting with tau441 to bind to MTs when expressed together in HEK293T cells. Data (n=6- 7/group) are shown as mean ± SEM. NTR = N-terminal projection region; PRR = proline-rich region; MTBR = microtubule-binding region; E4a = exon 4a; E6 = exon 6; ns = not significant; \**p*<0.05; \*\**p*<0.01; \*\*\**p*<0.001; \*\*\*\**p*<0.0001; unpaired t-test.

The experiments above suggest that big tau shows strong microtubule binding due to additional microtubule-binding sites. If true, then we would predict that big tau should compete with tau441 for microtubule binding. To test this, we co-expressed a different ratio of big tau and tau441 in HEK293T cells while maintaining the total level of tau expression consistent. When we performed the microtubule- fractionation analysis and subsequent western blots with these cells, we found that replacing tau441 with big tau results in an increased enrichment of big tau in the microtubule-bound fraction while drastically reducing the amount of tau441 from the same fraction (**Fig. 3f,g**). Notably, even substituting ∼20% of tau441 with big tau led to a robust enrichment of big tau in the microtubule-bound fraction (**Fig. 3f,g**). As such, these results collectively indicate that big tau can indeed outcompete tau441 for microtubule- binding *in vitro*, supporting the conclusion that big tau has a stronger binding affinity to microtubules compared to regular tau.

### Big tau is significantly less prone to aggregation

So far, we have shown that big tau has reduced hyperphosphorylation and increased turnover and microtubule binding. As all these mechanisms are implicated in tau pathology, we speculated that big tau could be more resistant to aggregation. We analyzed brain samples from aged mice using western blot analysis and an antibody that detects abnormal conformation resembling AD tau fibrils (i.e., MC-1). Here, we found that regular isoforms of endogenous mouse tau are subject to such conformational changes in an age-dependent manner, but not big tau (**Fig. 4a-c**). Inspired by this, we examined the aggregation propensity of big tau using an established cellular seeding assay^25^. Specifically, we expressed either FLAG-tagged big tau or tau441 (both containing the aggregation-promoting P301L mutation) in HEK293T cells and subsequently treated them with recombinant tau seeds to induce intracellular tau aggregation. Then, we harvested cells in a Triton-containing buffer and performed ultracentrifugation-based fractionation, resulting in Triton-soluble and insoluble fractions containing monomeric and aggregated forms of tau species, respectively. Western blot analysis using both the Dako-tau antibody and the FLAG antibody revealed that tau441 dramatically aggregated upon recombinant tau seed treatment, consistent with previous reports^25^ (**Fig. 4d-g**). In contrast, under the same tau seed treatment condition, big tau was dramatically resistant to seed-induced aggregation (**Fig. 4d-g**).

**Figure 4.**
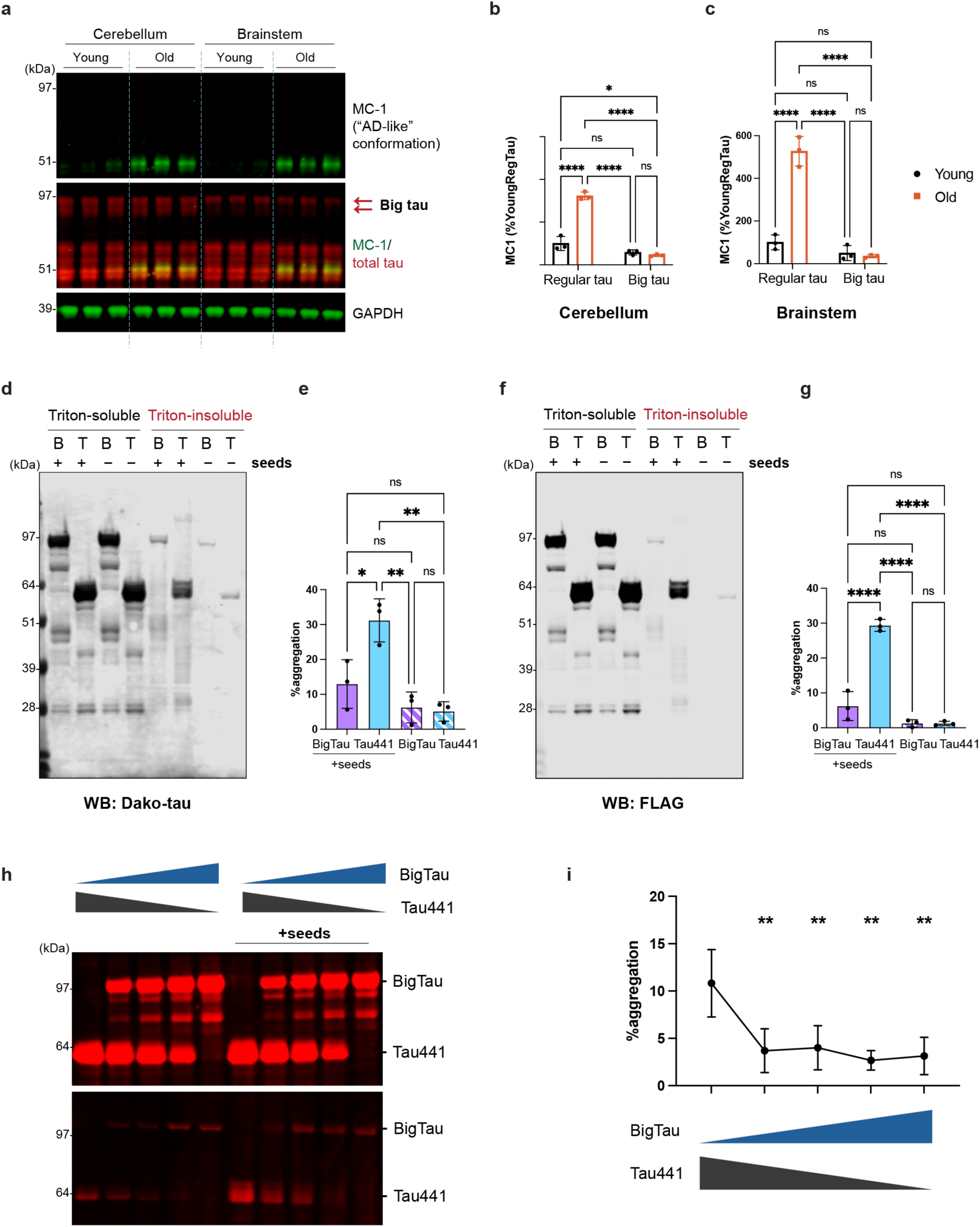
Big tau has a significantly decreased aggregation propensity compared to tau441. **a-c,** Representative western blots probed by the MC-1 antibody showing an age-dependent increase in abnormal conformational changes associated with endogenous mouse tau (**a**) and their quantification for cerebellum (**b**) and brainstem (**c**), both normalized to GAPDH. **d-e,** Representative western blots probed with the Dako-tau antibody (**d**) and their quantification (**e**) showing the degree of aggregation between big tau and tau441 upon the recombinant tau seed treatment. **f-g,** Representative western blots probed with the FLAG antibody (**f**) and their quantification (**g**) showing the degree of aggregation between big tau and tau441 upon the recombinant tau seed treatment. **h-i,** Representative western blots probed with the Dako-tau antibody (**h**) and their quantification (**i**) showing reduction in total tau aggregation upon replacing tau441 with big tau. Data (n=3-4/group) are shown as mean ± SEM. ns = not significant; \**p*<0.05; \*\**p*<0.01; \*\*\**p*<0.001; \*\*\*\**p*<0.0001; one-way ANOVA.

Based on the discovery that big tau has reduced aggregation capacity, we next tested if replacing tau441 with big tau while maintaining the total tau expression consistent can reduce total tau aggregation. Similar to our microtubule-binding competition assay for big tau and tau441, we expressed a different ratio of big tau and tau441 in HEK293T cells, induced tau aggregation using recombinant tau seeds, and performed detergent fractionation. Interestingly, we observed a decrease in total tau aggregation as we continued to substitute tau441 with big tau (**Fig. 4h,i**), which further demonstrates a significantly reduced aggregation propensity of big tau in this cellular tau aggregation assay.

### Big tau resists aggregation in vivo

To further determine big tau’s aggregation propensity in a mammalian model, we employed a previously published virus-mediated method to express tau in the mouse brain^26^. Specifically, we performed an intracerebroventricular injection into postnatal day 0 pups (P0-ICV) with adeno-associated viruses (AAVs) to overexpress either big tau or tau441 (both containing the aggregation-promoting P301L mutation) in the mouse brain (**Fig. 5a**). Expression of regular tau with the P301L mutation using this method has been reported to result in robust tau aggregation and hyperphosphorylation in the 6-mo mouse brain^26^. Consistent with this finding, when we performed ultracentrifugation-mediated fractionation to isolate the sarkosyl-insoluble fraction that includes aggregated tau species^26,27^, we found that tau441^P301L^ was robustly aggregated and hyperphosphorylated, as detected by the Dako-tau antibody and a phospho-tau antibody (i.e., PHF1) in western blots (**Fig. 5b,c**). In contrast, we found that big tau is strikingly less prone to these pathological changes despite including the same pathogenic tau mutation, as tau441 was approximately 15-fold more likely to become aggregated and hyperphosphorylated in the brain of this mouse model (**Fig. 5b,c**). Of note, AAVs were injected to express similar amounts of tau441 and big tau between two groups as demonstrated by qPCR analysis (**Extended Data** Fig. 6a,b), indicating that the observed pathology-resisting properties of big tau are not due to the expression level differences, but due to the big tau protein itself. Moreover, differences between big tau and tau441 in their pathological changes were still present even in older mice (9-mo) (**Fig. 5d,e**), further demonstrating that big tau can uniquely evade pathological modification such as aggregation and hyperphosphorylation.

**Figure 5.**
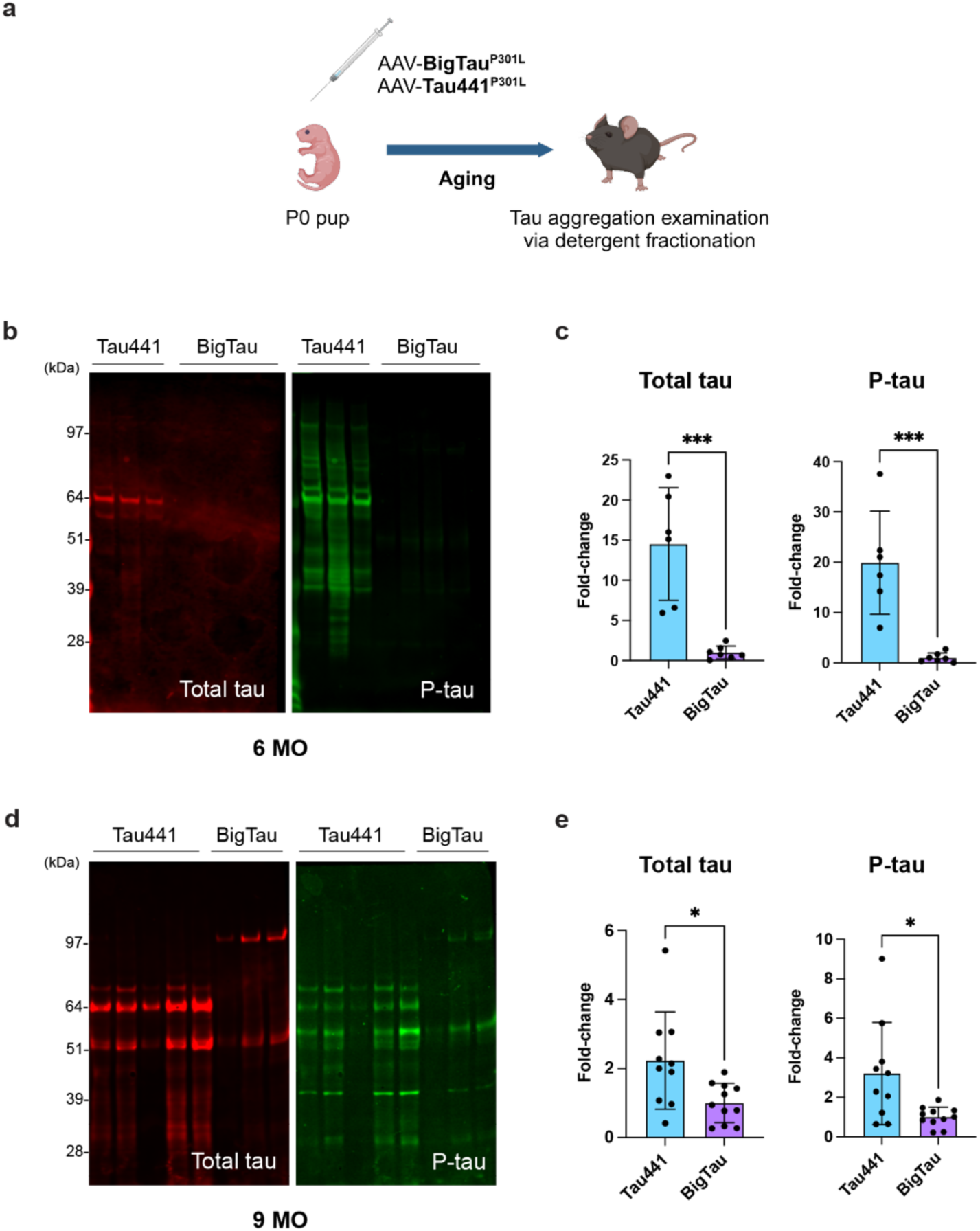
Big tau does not become pathologically aggregated and hyperphosphorylated in the AAV-based mouse model. **a,** A schematic diagram depicting the mouse model established via the intracerebroventricular injection into postnatal day 0 pups (P0-ICV) of adeno-associated virus (AAV) expressing either big tau or tau441 carrying the P301L aggregation-promoting mutation. **b-c,** Representative western blots probed with the Dako-tau (total tau) antibody and PHF1 (phospho-tau; p- tau) antibody (**b**) and their quantification (**c**) of the sarkosyl-insoluble fraction extracted from 6-months- old (6MO) AAV mice. **d-e,** Representative western blots probed with the Dako-tau (total tau) antibody and PHF1 (phospho-tau; p-tau) antibody (**d**) and their quantification (**e**) of the sarkosyl-insoluble fraction extracted from 9-months-old (9MO) AAV mice. Data (n=6-10/group) are shown as mean ± SEM. \**p*<0.05; \*\*\**p*<0.001; unpaired t-test.

### AD patients have a significantly higher level of big tau in the cerebellum

Previously, big tau has not been discussed in the context of the human brain. We analyzed big tau transcript levels in human brains using data from the Mayo RNAseq study^28^ and found that the big tau transcript level is significantly higher in the cerebellum than in the temporal cortex in humans (**Fig. 6a**), consistent with our earlier observation in mice (**Fig. 1e-f**). Interestingly, we found an additional difference in big tau levels per disease status. Compared to cognitively normal control individuals, AD patients displayed a significantly higher level of big tau transcript (normalized to the level of total tau transcript) in the cerebellum (**Fig. 6b**), a brain region lacking overt tau pathology. In contrast, AD patients and controls showed no significant difference in the big tau transcript level in the temporal cortex, one of the brain regions that are heavily impacted by tau pathology during the disease course (**Fig. 6c**). Of note, we did not find a sex-difference effect in the big tau level (**Extended Data** Fig. 7a**- d**).

**Figure 6.**
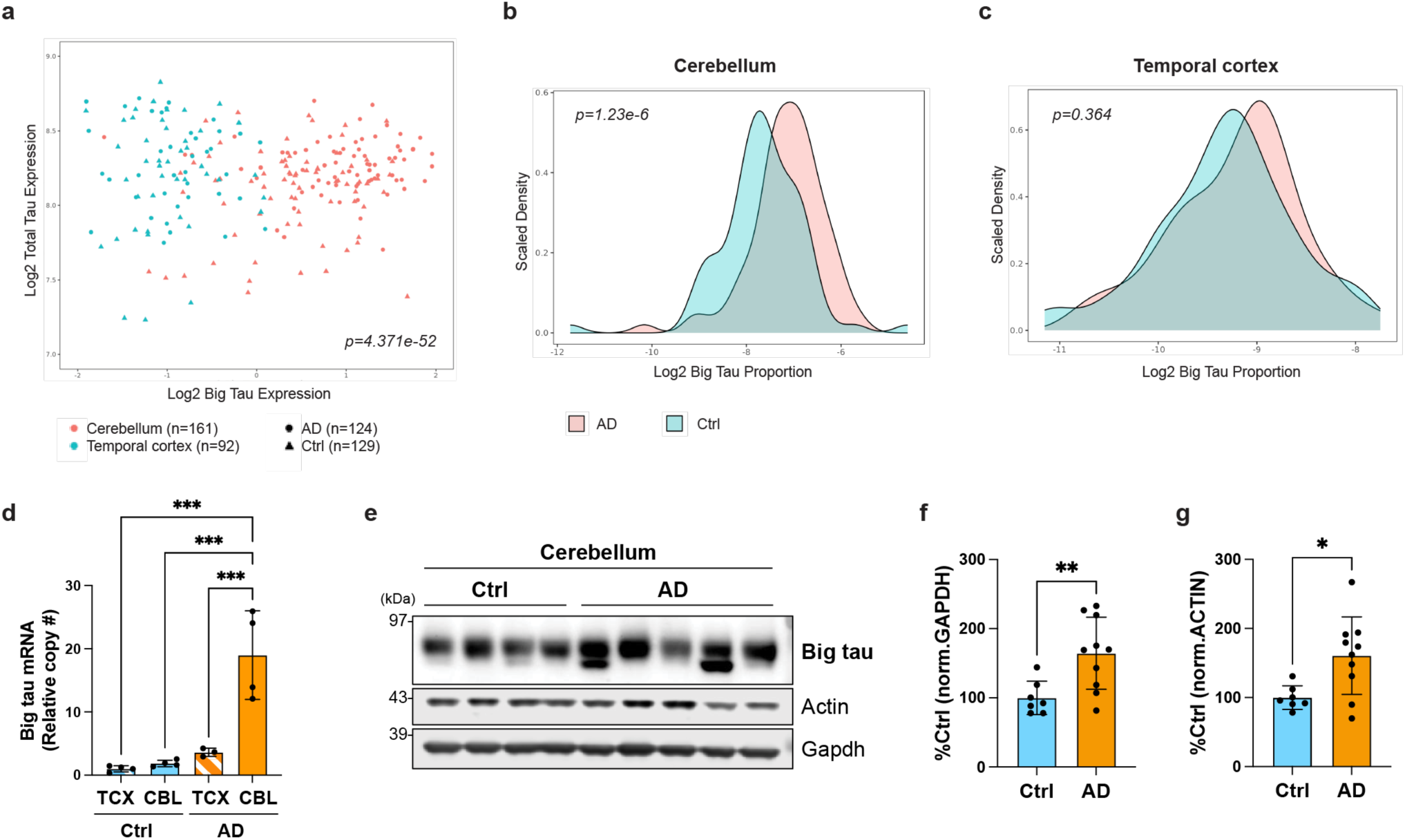
The cerebellar big tau level is significantly higher in AD patients than in control individuals. **a,** Analysis of the big tau transcript level across cerebellar and temporal cortical tissues from AD patients (n=124) and control individuals (Ctrl; n=129). **b-c,** Comparison of the big tau transcript level (normalized to the total tau level) in the cerebellum (**b**) and the temporal cortex (**c**) between AD patients and controls. **d,** A graph showing the relative copy number of the big tau transcript measured by digital PCR in cortical and cerebellar tissues of AD patients and controls. **e-g,** Representative western blots (**e**) and their quantification normalized to GAPDH (**f**) or actin (**g**) showing a significantly higher level of cerebellar big tau protein in AD patients compared to controls. Data (n=3- 4/group for digital PCR; n=7-10/group for western blot) are shown as mean ± SEM. CBL = cerebellum; TCX = temporal cortex; \**p*<0.05; \*\**p*<0.01; \*\*\**p*<0.001; one-way ANOVA for digital PCR; unpaired t- test for western blots.

Based on the RNAseq analysis results, we further examined this phenomenon by performing digital PCR to measure the absolute amount of big tau transcript in postmortem brain samples of AD patients and control individuals (**Extended Data Table 1**). Here, we found that the amount of big tau transcript is significantly higher in the AD cerebellum compared to the control cerebellum, control cortex, or AD cortex (**Fig. 6d**). We also performed western blots using our human big tau-specific antibody (**Extended Data** Fig. 2a,b**,d**) to detect big tau protein in the lysates of cerebellar tissues of AD patients and control individuals (**Extended Data Table 1**). Consistent with the RNAseq analysis and digital PCR results, we found that the cerebellar big tau protein level is significantly higher in AD patients (**Fig. 6e-g**). Our analyses using both mouse and human brain samples collectively indicate that big tau is robustly expressed in the cerebellum, which was rather surprising given the prior belief that big tau exists predominantly in the PNS. Furthermore, our data demonstrate that AD patients show a higher level of pathology-resisting big tau in the brain region protected from developing tau pathology in AD, suggesting an intriguing link between the elevated level of pathology-resisting big tau and the absence of AD-tau pathology.

## Discussion

Every neurodegenerative disorder is characterized by selective vulnerability, where certain cell populations and brain regions are predominantly affected, while others are seemingly resistant to the development of pathology and its associated toxicity^1,29–34^. Understanding the regulators of this vulnerability remains an important outstanding question in the neurodegeneration field, AD being no exception. Despite the universal expression of tau in the brain, tau pathology only affects selective brain regions in AD. In this study, we demonstrated that the understudied big tau isoform, highly expressed in the brain regions that do not develop severe tau pathology in AD, possesses several biochemical properties that are likely to render it resistant to various pathological changes (**Extended Data** Fig. 8). Interestingly, we found that AD patients display a significantly higher level of cerebellar big tau compared to control individuals, which suggests a potential causal link between an increased level of pathology-resisting big tau and the absence of tau pathology in the AD cerebellum.

So far, many tau isoform studies have focused on the 3R versus 4R aspect of tau, given the significance of the 3R:4R tau isoform ratio in tauopathies and the observation that certain tau mutations can shift tau splicing to promote the expression of 4R tau^13^. In contrast, big tau has been rarely studied in the brain despite its initial discovery in the 1990s. In the central nervous system (CNS), big tau has been known to be mostly expressed in cell types whose processes extend to the periphery, such as dorsal root ganglion, superior posterior ganglion, and spinal motor neurons^35,36^. However, some cell types whose processes are contained within the CNS such as retinal ganglion cells and optic nerve were also reported to express big tau^36,37^. The potential presence of big tau transcript has been described in the rat cerebellum^36^ but it was not extensively characterized. As such, our findings of region-specific expression of big tau in the mouse and human brain are particularly interesting as they expand our current understanding of tau biology. Importantly, to our knowledge, our study is the first study to examine the expression levels of big tau in the human brain.

We discovered big tau’s significantly diminished aggregation propensity as one of the key differences between big tau and regular tau isoforms. Consistent with our findings, AD and tauopathy patients typically do not develop notable peripheral tau pathology^38,39^, further supporting the notion that big tau resists pathological aggregation. Several factors may contribute to the suppression of big tau aggregation, such as the sheer length of big tau’s NTR. It is well-known that the “K18” tau fragment, comprised exclusively of the C-terminal repeat domain, aggregates much faster than the full-length tau^40^, and it has been reported that the N-terminal tau fragment can inhibit the aggregation of the full- length tau^41^. These findings raise the possibility that the expanded disordered NTR lowers tau’s aggregation propensity. Big tau has an NTR three times larger than any other tau isoforms, suggesting that this “buffering” function of NTR could be enhanced. Also, as evidenced by a pioneering cryo-EM study using tau fibrils isolated from AD brains, tau’s N-terminal amino acid residues (^7^EFE^9^) are suggested to interact with the core of tau fibrils to stabilize the whole fibrillar structure^42^. Hence, it is possible that the longer NTR of big tau disfavors the EFE motif from looping back and interacting with the fibril core, ultimately making tau fibrils less stable. Potentially supporting this idea is our finding that the MC-1 antibody, which recognizes an epitope formed by the EFE motif and the tau fibril core^42–44^, fails to detect big tau.

Another factor that can potentially contribute to big tau’s reduced aggregation is its enhanced microtubule-binding capacity. Once detached from microtubules, tau is believed to become more susceptible to pathological aggregation and accumulation, along with abnormal changes in cytoskeletal dynamics in the nervous system^45^. Indeed, microtubule defects are pronounced in AD and other tauopathies^46,47^, and multiple pathogenic tau mutations are reported to decrease tau’s affinity to microtubules and impact its ability to promote microtubule assembly^48^. Based on our findings, big tau has a lower chance of being detached from microtubules, hereby limiting its propensity to aggregate and preventing downstream pathological changes to the neuronal cytoskeleton. Besides, we also show evidence that big tau displays less hyperphosphorylation—a key hallmark of tau pathology that ultimately promotes tau aggregation^19^. Moreover, big tau is more readily ubiquitinated *and* degraded, being less prone to accumulate pathological modifications by serving as a better substrate for the cell’s degradation machinery. Combined, all these mechanisms likely contribute to big tau’s resistance to aggregation.

In addition to the properties discussed above, isoforms of the same protein can critically diversify the biological functions of one gene via their distinct protein structures^49^, which could also be the case for big tau. While we still do not know if big tau possesses unique physiological roles different from other major tau isoforms, some roles for big tau have been proposed such as more efficient organelle transport in the axon or maintenance of increased spacings between microtubules where they are needed^14^. Also, our work uncovered that big tau’s NTR can strengthen microtubule-binding activities, which was surprising as the NTR of tau has not been previously thought to bind to microtubules unlike MTBR or PRR^21–23^. As such, future studies are warranted to investigate big tau- specific biological functions using a novel platform such as mouse models genetically engineered to be depleted of big tau.

Taken together, our data clearly demonstrate that big tau is drastically distinct from other tau isoforms, highlighting the importance of investigating neurodegenerative disease proteins in an isoform- specific manner. Apolipoprotein E (apoE) isoforms and their critical implication in distinctly modulating the risk of sporadic AD are classic examples of isoform-specific effects on the disease progression^50^. The potential involvement of α-synuclein isoforms in different pathophysiological consequences has also been discussed for Lewy body dementia^51^. Recently, another study discovered a novel tau isoform (“W-tau”) with a truncated protein structure generated by *MAPT* intron 12 retention, which also possesses a reduced aggregation propensity similar to big tau^52^. As seen by these examples, additional investigations on less-studied tau isoforms may provide us with critical insights into tau biology and novel ideas for developing potential therapies.

Can we potentially leverage pathology-resisting properties of big tau to develop tau-targeting therapeutic strategies for AD and other tauopathies in the future? Our data from cellular assays have shown that substituting regular tau (tau441) with big tau while maintaining the total tau amount the same may be beneficial since big tau can (1) bind to microtubules more strongly, and (2) suppress the total tau aggregation. Once big tau is confirmed to be a safe alternative to regular tau in the adult brain, switching CNS-tau isoforms (at least partially) to big tau using anti-sense oligonucleotides (ASOs)^53^ will become an interesting therapeutic strategy. To explore this possibility, it would be important to understand this region-specific splicing of big tau in the brain. Of note, *MAPT* exon 4a (753 nucleotides) is considered a “large exon” as most vertebrate exons do not exceed 400 nucleotides^54^. The longer exon length can prevent the spliceosome from recognizing the exon to splice^54,55^, which may partially explain the default exclusion of big tau exons during the splicing events in certain brain regions. Identifying splicing factors responsible for the regulation of big tau-specific exons of *MAPT* may help us elucidate the brain region- specific big tau splicing mechanisms.

In sum, our study has demonstrated that big tau can uniquely resist pathological changes related to AD. Intriguingly, the levels of big tau are elevated in the brain regions that typically do not develop tau pathology in AD, suggesting that alternative splicing of *MAPT* that leads to the expression of big tau may explain the regional brain resistance to tau pathology in AD. As the first study to systematically explore big tau biology, our work expands our understanding of the complexity underlying tau pathology and provides uncharted paths for potential therapeutic approaches for AD and related tauopathies.

## ACKNOWLEDGEMENTS

The authors would like to thank Sameer Bajikar, Hamin Lee, Ji-yoen Kim, Youngdoo Kim, and Jian Zhou for their feedback on this manuscript and other Zoghbi lab members for their insightful discussion. The authors would like to further thank the Baylor College of Medicine (BCM) Gene Vector Core. The authors deeply appreciate patients and their families for their generous donation of brain samples for the study. This study was supported by the following funding sources: Baylor College of Medicine Center for Alzheimer’s and Neurodegenerative Diseases (CAND) Scholars Program (D.C.C.); BrightFocus Foundation Postdoctoral Fellowship Program in Alzheimer’s Disease Research A2023002F (D.C.C.); United States Department of Agriculture (USDA/ARS) under Cooperative Agreement No. 58- 3092-0-001 (H.K.Y.); Duncan NRI Zoghbi Scholar Award (H.K.Y.); JPB Foundation MR-2023-4260 (H.Y.Z., B.T.H.); Howard Hughes Medical Institute (H.Y.Z.); Massachusetts Alzheimer’s Disease Research Center (NIA P30AG062421); The Association for Frontotemporal Degeneration (AFTD) 2023-001 (S.B.); Cancer Prevention and Research Institute of Texas (CPRIT) RR220094 (S.B.); Eunice Kennedy Shriver National Institute of Child Health & Human Development; Baylor College of Medicine Intellectual and Developmental Disabilities Research Center (BCM-IDDRC) (NICHD P50HD103555)

## AUTHOR CONTRIBUTIONS

D.C.C., S.B., and H.Y.Z conceptualized the study. D.C.C., X.D., J.P.R., S.B., and H.Y.Z. designed the experiments and interpreted the data. D.C.C., X.D., J.P.R., A.L.H, B.T., R.R., and F.A.N. performed the experiments. H.K.Y. and M.D. analyzed the human RNAseq database. B.T.H. provided the human postmortem brain tissues. D.C.C. and H.Y.Z. obtained funding for the study. D.C.C. wrote the original draft of the manuscript. D.C.C., X.D., H.K.Y., J.P.R., B.T., M.D., S.B., and H.Y.Z. edited the manuscript. All authors discussed the results and commented on the manuscript.

## CORRESPONDING AUTHOR

Correspondence to Dr. Huda Y. Zoghbi.

## METHODS

### Antibodies

We generated rabbit polyclonal antibodies specific for mouse big tau protein (using peptides QASQPEGPGTGPMEEGHE) and human big tau protein (using peptides EEVDEDRDVDESSPQDSPPS) in collaboration with Pacific Immunology (Ramona, CA, USA). Tau antibodies such as PHF1 (pS396/S404 of tau) and MC-1 (conformational epitope of tau) were kind gifts from the late Dr. Peter Davies (Feinstein Institute for Medical Research, North Shore LIJ Health Care System). The following antibodies were purchased from the manufacturers described: mouse anti-tau antibody (tau-5; Cat# ab80579) and rabbit anti-GAPDH antibody (Cat# ab181602) from Abcam (Cambridge, United Kingdom), rabbit anti-tau polyclonal antibody (Cat# A0024) from Dako-Agilent Technologies (Glostrup, Denmark; Santa Clara, CA), mouse anti-phospho-tau (pS202/T205) antibody (AT8; Cat# MN1020) from Thermo Fisher Scientific (Waltham, MA, USA), mouse anti-FLAG (M2) antibody (Cat# F3156) and mouse anti-α-tubulin (Cat# T6199) from Sigma-Aldrich (St. Louis, MO, USA), mouse anti-GAPDH (Cat# 2-RGM2) from Advanced ImmunoChemical (Long Beach, CA, USA), mouse anti-β-actin (Cat# 3700S), rabbit anti-HA (Cat# 3724S), and rabbit anti-PPIA (Cat# 2175S) from Cell Signaling Technology (Danvers, MA, USA), HRP-conjugated donkey anti-mouse secondary antibody (Cat# 715-035-150) from Jackson ImmunoResearch Laboratories (West Grove, PA, USA), HRP-conjugated anti-rabbit secondary antibody (Cat# 170-5046) from Bio-Rad (Hercules, CA, USA), and goat anti-Mouse and anti-rabbit secondary antibodies conjugated with Alexa Fluor (Cat# A32723, A32740) from Thermo Fisher Scientific.

### Chemicals and reagents

The following chemicals and reagents were purchased from the manufacturers described: isopropyl β-D-1-thiogalactopyranoside (IPTG) from Gold Biotechnology (St. Louis, MO, USA), L-arabinose, dithiothreitol (DTT), dextran sulfate, cycloheximide, guanosine 5’-triphosphate sodium salt hydrate (GTP), paclitaxel (Taxol), N-lauroylsarcosine sodium salt (sarkosyl), and thioflavin S from Sigma-Aldrich, EGTA from bioWORLD (Dublin, OH, USA), EDTA, imidazole, and Pierce Universal Nuclease from Thermo Fisher Scientific, Xpert Phosphatase Inhibitor Cocktail Solution and Xpert Protease Inhibitor Cocktail Solution from GenDEPOT (Katy, TX), pre-formed microtubules from Cytoskeleton (Denver, CO, USA), and MG132 from EMD Millipore (Burlington, MA, USA).

### Cell culture and transfection

HEK293T cells and N1E-115 cells (obtained and certified from ATCC) were maintained in the DMEM (Thermo Fisher Scientific, Cat# 10-013-CV) supplemented with 10% fetal bovine serum (Bio-Techne [Minneapolis, MN, USA], Cat# S11150) and 1% Antibiotic- Antimycotic (Thermo Fisher Scientific, Cat# 15240096). Cells were passaged every 3-4 days at 90% confluence. For transient plasmid transfection, following the manufacturer’s instructions, cells were transfected with plasmids encoding cDNA of interest or siRNAs targeting genes of interest combined with the Lipofectamine™ 2000 transfection reagent (Thermo Fisher Scientific) or Lipofectamine™ RNAiMAX transfection reagent (Thermo Fisher Scientific), respectively, in serum-free Opti-MEM (Thermo Fisher Scientific). Cells were transfected for 48-74 hrs and harvested for further analyses.

### Mouse studies

To examine endogenous mouse tau isoforms, brain tissues of C57/B6 (Strain# 000664) and FVB (Strain # 001800) mice from the Jackson Laboratory (Bar Harbor, ME, USA) were used. To establish the tau-AAV mouse model, intracerebroventricular injections of AAV were performed using C57/B6 mouse pups at postnatal day 0 (P0 ICV injection) modifying previous protocols^26,56^. In brief, newborn mice were cryoanesthetized and placed on a cold metal plate. To target lateral ventricles, the skull was pierced using a 30-gauge needle (posterior to bregma and 2 mm lateral to the midline) and AAV8 expressing Tau441^P301L^ and BigTau^P301L^ was injected (2E+10 viral particles/ventricle; 2 µl/ventricle). Pups were placed on the warming pad post-injection and returned to their home cage after beginning to move actively. All mice were housed (up to 5 mice per cage) with 14 hr light and 10 hr dark cycle at RT, provided with water and standard rodent diet *ad libitum*. All mouse procedures were approved by the Baylor College of Medicine Institutional Animal Care and Use Committee (IACUC) and are in accordance with the National Institutes of Health Guide for the Care and Use of Laboratory Animals (NIH Publications No. 80-23, revised 1996).

### Sample preparation and western blots

For cells, samples were harvested and lysed in the lysis buffer (50 mM Tris HCl [pH 7.4], 274 mM NaCl, 5 mM KCl, 5 mM EDTA, 1% Triton-X-100, 1 mM PMSF, protease and phosphatase inhibitor cocktails). For the mouse brains, at the time of harvest, mice were deeply anesthetized with a mix of ketamine 37.5 mg/ml, xylazine 1.9 mg/ml, and acepromazine 0.37 mg/ml and underwent transcardial perfusion with 20 ml of ice-cold PBS. The brains were collected, dissected along the midline, immediately frozen, and stored at -80 °C until lysed in the lysis buffer described above for biochemical analysis.

Cell or brain tissue lysates were centrifuged at 16,000 x g for 15-20 min at 4 °C, and the supernatant was collected for standard BCA protein assay (Thermo Fisher Scientific). Appropriate volumes of lysate (1-2 μg of protein per μl) in 1x SDS sample buffer (Thermo Fisher Scientific) were heat-denatured 95 °C. Samples (10-20 ug of protein per well) were run on precast 4-12% Bis-Tris gels with the MOPS running buffer (Thermo Fisher Scientific) and transferred onto nitrocellulose membranes (Bio-Rad) in the Tris-glycine transfer buffer supplemented with 10% methanol. Membranes were blocked in 5% non-fat dry milk in TBS/0.1% Triton-X-100 (TBST) for 30 min to 1 hr at RT, and incubated with primary antibody rocking overnight at 4 °C. The following antibodies were used at the indicated dilution: tau-5 (1:2,000), Dako anti-tau (1:8,000), AT8 (1:1,000), MC-1 (1:1,000), PHF1 (1:1,000), anti-FLAG (1:5,000), anti-HA (1:5,000), anti-tubulin (1:1,000), anti-GAPDH (1:10,000), anti-actin (1:1,000), anti-PPIA (1:1,000), mouse big tau (1:1,000), and human big tau (1:500). The membranes were washed with TBST three times and subsequently incubated with secondary antibodies for 1 hr at RT. For HRP-conjugated secondary antibodies, blots were detected using the Pierce ECL detection kit (Thermo Fisher Scientific) and imaged using Amersham™ Imager 680 (GE Healthcare, Chicago, IL, USA). For secondary antibodies conjugated with Alexa Fluor, blots were imaged with the Odyssey CLx imager (LI-COR Biosciences, Lincoln, NE, USA). Protein bands were quantified using the Image Studio software (Li-COR Biosciences).

### Big tau ubiquitination assay

FLAG-tagged big tau or tau441 was co-expressed with HA-tagged ubiquitin in HEK293T cells for 48 hrs. Before harvest, cells were treated with MG132 for 6 hrs to block the proteasomal degradation. After cell lysis, anti-FLAG (M2) Magnetic beads (Sigma-Aldrich) were used to immunoprecipitate big tau or tau441 from the cell lysates following the manufacturer’s instructions. Western blot was subsequently performed using the HA antibody to examine the ubiquitination of big tau or tau441.

### Cycloheximide chase assay

HEK293T cells were transfected with either big tau or tau441 (both FLAG-tagged) for 48 hrs. Subsequently, cell media was supplemented with cycloheximide at the final concentration of 50-100 μg/ml. Cells were collected at desired time points and kept frozen at -20°C until further western blot analysis.

### Cell-based microtubule binding assay

To compare the ability of tau441 and big tau to bind microtubules (MTs) in cells, HEK293T cells were transfected with the corresponding plasmid for 24 hrs. By adapting previous protocols^57,58^, cells were lysed in 200 μl of PEM buffer (80 mM PIPES, pH 6.8, 1 mM EGTA, 1 mM MgCl_2_) supplemented with 0.1% Triton X-100, 2 mM GTP, 20 µM Paclitaxel, and a mix of protease and phosphatase inhibitors at 37 °C for 30 min and then centrifuged at 100,000 × g for 1 hr at 37 °C. The supernatant was collected as an unbound fraction. The pellet (MT-bound fraction with MT-bound proteins) was washed twice, resuspended in 200 μl of PEM buffer, and sonicated. LDS loading buffer was added to both fractions and equal amounts of each fraction were analyzed by Western blot as described above. The percentage of MT binding was calculated as (protein in the MT-bound fraction) / [(protein in the MT-bound fraction) + (protein in the unbound fraction)].

### Peptide array overlay assay

To map the epitopes within big tau that bind to microtubules, human big tau protein was synthesized as fragments of linear peptides (with the acetylated N-terminus) covalently bound to a cellulose membrane by the C-terminus. These membranes were produced using the SPOT synthesis technique^59^ by JPT Peptide Technologies (Berlin, Germany). Each “spot” on the membrane contained approximately 5 nmol of 15-mer peptide fragments tiling the entire sequence of human big tau while overlapping with the next peptide fragment by 11 amino acid residues. Spots were arrayed in a grid with 0.37 cm spacing from one another, up to 20 spots per row. For the overlay assay, the membrane was incubated in 10% non-fat dry milk in TBS overnight at 4 °C. Next day, the membrane was washed with TBS three times and subsequently incubated with pre-formed, Taxol-stabilized microtubules (Cytoskeleton, Cat# MT002) at a final concentration of 2.5 μg/ml. After three washes with TBS, the membrane was incubated with the anti-tubulin antibody followed by an HRP-conjugated secondary anti-mouse antibody to reveal microtubule binding via classical western blot.

### Cellular (big)tau aggregation assay

To isolate the triton-soluble and insoluble fractions from cells expressing tau based on previous protocols^60,61^, cells were transfected to overexpress human big tau or tau441 containing the aggregation-promoting P301L mutation and treated with recombinant tau seeds 6 hrs after transfection. At 48 hrs post-transfection, cells were harvested and lysed in the lysis buffer described above and ultracentrifuged at 100,000 x g for 30 min at 4 °C using the TLA-110 rotor and the Optima™ MAX-XP tabletop ultracentrifuge (Beckman, Brea, CA). The supernatants were collected as “triton-soluble fraction” and the pellets were washed with the lysis buffer. The resulting pellets were resuspended in the lysis buffer supplemented with SDS at 1% final concentration and sonicated as “triton-insoluble fraction.” Each fraction was subject to western blot analysis.

### Detergent fractionation of mouse brain tissues

To isolate sarkosyl-soluble and insoluble fractions from mouse brain tissues following the previous protocols^56,61^, mouse brain tissues were homogenized in 10x volume of Buffer A (10 mM Tris-HCl, 80 mM NaCl, 1 mM MgCl_2_, 1 mM EGTA, 0.1 mM EDTA, 100 mM DTT, 1 mM PMSF, protease and phosphatase inhibitor cocktails) and ultracentrifuged at 150,000 x g for 70 min at 4 °C using the TLA-110 rotor and the Optima™ MAX-XP tabletop ultracentrifuge. Supernatant was kept as the “soluble fraction.” The pellet was resuspended in Buffer B (10 mM Tris-HCl, 850 mM NaCl, 1 mM EGTA, 10% sucrose) and centrifuged at 150,000 x g for 15 min at 4 °C to remove debris. The pellet was kept for potential future analyses, and the supernatant was collected to be incubated with 1% sarkosyl for 1 h at RT. After incubation, the sample was ultracentrifuged again at 150,000 x g for 40 min at 4 °C using the TLA-110 rotor. The supernatant was collected as the “sarkosyl-soluble fraction”. The pellet was washed with and resuspended in ice-cold PBS and sonicated as the “sarkosyl-insoluble fraction.”

### Quantitative RT-PCR (qPCR) and digital PCR

Total RNA was isolated from mouse or human brain tissues using TRIzol™ reagents (Thermo Fisher Scientific, Cat# 15596026) following the manufacturer’s instructions and appropriately stored at -80°C until further use. To synthesize cDNA, reverse transcription (RT) was performed with 500-1000 ng of RNA at 37 °C for 15 min followed by heat inactivation at 85 °C for 5 sec using the PrimeScript™ RT Reagent Kit (Takara Bio, Kusatsu, Shiga, Japan) following the manufacturer’s instructions.

For qPCR, 25 ng of cDNA was mixed with PowerUp™ SYBR™ Green Master Mix (Thermo Fisher Scientific), gene-specific primers, and water in a total volume of 12.5 µl. All qPCR mixed reactions were run in duplicates on the C1000TM Thermal Cycler (Bio-Rad). Relative gene expression was normalized to control (GAPDH) and calculated using the regular 2^-ΔΔCt^ method. The following gene-specific primers were ordered from Sigma-Genosys (Spring, TX, USA): total mouse tau (mMapt; Forward: 5’-GAGGGAATAAGAAGATTGAAACCC-3’; Reverse: 5’- CAATTTCTGCTCCATGGTCTG-3’), mouse big tau (Forward: 5’-ACGGAGATCCCAGAAGGAAT-3’; Reverse: 5’GAGCCCCAGACATGCTAGAC-3’), mGapdh (Forward: 5’- AGGTCGGTGTGAACGGATTTG-3’; Reverse: 5’-TGTAGACCATGTAGTTGAGGTCA-3’), total human tau (hMAPT; Forward: 5’-CCAGTCCAAGTGTGGCTCAAAG-3’; Reverse: 5’- GCCTAATGAGCCACACTTGGAG-3’), human big tau (Forward: 5’- CAAGGAGGAGGTGGATGAAGACC-3’; Reverse: 5’-TTGGAGAGGAAATCCACAGGGAG-3’).

For digital PCR, 30 ng of cDNA was mixed with the ddPCR Supermix for Probes (Bio-Rad) and the reactions were run on the QX200 Droplet Digital PCR System platform (Bio-Rad). The expected detection targets consisted of big tau mRNAs (HEX fluorophore) and TUBB3 mRNAs (FAM fluorophore). PCR cycling consisted of a denaturation step at 95 °C for 30 s followed by annealing/extension at 60 °C for 1 min with a ramp at 2 °C per sec (40 cycles), and a final extension step at 60 °C for 5 min. Data were analyzed using QuantaSoft Analysis Pro software (Bio-Rad). The following digital PCR primers and probes were ordered from Integrated DNA Technologies IDT (Coralville, IA, USA): human big tau (primer 1: 5’-AGAGATCCCAGCCTCAGAGC -3’; primer 2: 5’-CCCGTCTTTGCTTTTACTGA -3’; probe: 5’-CCTCTGAAAAGCAGCCT-3’); hTUBB3 (primer 1: 5’- GGCCTTTGGACATCTCTTCAG-3’; primer 2: 5’-CCTCCGTGTAGTGACCCTT-3’; probe: 5’- CGGCCCCACTCTGACCAAAGAT-3’).

### Recombinant tau seed production

Human cDNA encoding the K18 tau fragment (a truncated form of tau consisting of only the four microtubule-binding repeats) was subcloned into the pET-28a bacterial expression vector with the His tag and transformed in BL21(DE3) competent cells (New England Biolabs). Cultures were grown to saturation overnight in Luria Broth on a shaker (220 rpm, 37 °C) and subsequently used to inoculate larger cultures the next morning (1:100 dilution). After the A_600_ reached 0.5 absorbance units, the protein expression of K18 tau was induced using a final concentration of 0.2% L-arabinose and 0.5 mM IPTG for 3.5 hrs. Cells were then collected by centrifugation and the pellets were stored at -80 °C until further use.

To purify recombinant tau K18 fragment, the frozen cell pellets were resuspended in the lysis buffer (10 mM HEPES [pH 7.4], 50 mM NaCl, 1 mM MgCl_2_, 0.5 mM DTT, supplemented with nuclease and proteinase and phosphatase inhibitors) and lysed with three freeze-thaw cycles in liquid nitrogen and the tepid water bath (25 °C). After removing debris by centrifuge, the supernatants were collected and added with NaCl at the final concentration of 500 mM. Tau K18 was affinity-purified against its His tag using TALON Metal affinity Resin (Takara bio) and eluted from the resin using a high concentration of imidazole (final concentration of 300 mM), following the manufacturer’s instructions. The elution fractions were pooled and dialyzed against 10 mM HEPES buffer (pH 7.4) overnight at 4 °C. After dialysis, the elution fraction was concentrated using speed vac.

To prepare the recombinant tau seeds, purified K18 tau was first diluted in the assembly buffer (10 mM HEPES [pH 7.4], 30 mM NaCl, 0.9mN MgCl_2_, 0.3 mM DTT, 6 μM EDTA, 1.5 μM EGTA) to the final concentration of 9 μM. Tau polymerization was induced by adding dextran sulfate at 0.04 μg/μl to the reaction and incubating the reaction at 37 °C overnight with agitation. The degree of polymerization was assessed by adding 7.5 μM of thioflavin S to an aliquot of reaction and measuring the thioflavin S binding 30 min later (excitation wavelength = 400 nm, emission spectrum = 460-600 nm). After confirming polymerization, the reaction was ultracentrifuged at 150,000 x g for 40 min at 4 °C using the TLA-110 rotor and the Optima™ MAX-XP tabletop ultracentrifuge. The resulting pellet was resuspended in ice-cold PBS and its concentration was measured via BCA protein assay. Before usage, the seeds were sonicated using Bioruptor Pico^®^ sonication device (Diagenode, Denville, NJ).

### Viral vector construct generation and virus production

The human big tau construct was generated by Gibson assembly of gBlocks gene fragments custom-ordered from IDT and cloned into the pcDNA3.1 vector that adds a FLAG tag at the end of the gene inserted. The sequence was verified by Sanger sequencing from Genewiz (South Plainfield, NJ). This big tau plasmid and the tau441 plasmid (both including the P301L mutation) were cloned into an AAV8 vector that includes the chicken β-actin promoter, a woodchuck post-transcriptional regulatory element (WPRE), and the bovine growth hormone polyA. Using these plasmids, AAVs were produced by the BCM Gene Vector Core by transfecting HEK293T cells with AAV helper plasmids for 96 hrs using iMFectin (GenDepot) and purifying AAVs by iodixanol density gradient. Virus titers were determined using WPRE-specific primers.

### Postmortem human brain tissues

All human postmortem brain tissues used for digital PCR and western blots in this study were provided by the Massachusetts Alzheimer’s Disease Research Center. Specifically, frozen tissues from the middle temporal cortex and cerebellum of AD patients (n=11) and control individuals (n=8) were obtained (Extended Data Table 1). Tissue donor anonymity was assured by the Massachusetts Alzheimer’s Disease Research Center.

### RNAseq database analysis

Aligned RNA-Seq data (in BAM format) and corresponding diagnostic information from postmortem brains of AD patients and controls were obtained from the AD Knowledge Portal^28^, which uses the Synapse platform for data hosting (syn5550404). The samples included tissues from both the cerebellum and temporal cortex. Read counts for exon 4a were used to quantify Big Tau isoform expression, while 3’UTR read counts were used to quantify total Tau (MAPT) expression. A total of 253 samples were downloaded, with 161 samples from the cerebellum and 92 samples from the temporal cortex. Raw read counts were extracted using featureCounts v2.0.0^62^, and total reads per sample were determined using samtools v1.10^63^. These raw counts were then normalized to the total reads per sample. Log2-transformed values of total Tau and Big Tau expression were visualized on a scatterplot, with points colored by tissue type and shaped by AD status (patient or control), implemented in R^64^. Big Tau proportions were calculated as the ratio of Big Tau to total Tau expression. Density plots of log2-transformed Big Tau proportions were generated separately for AD and control groups within each tissue type. The Kolmogorov-Smirnov test was used to evaluate the statistical significance of differences between AD and control groups, as well as between the cerebellum and temporal cortex.

### Data analyses

Statistical parameters such as the n number, measures (mean ± SEM), and statistical significance are reported in the figure Legends. GraphPad Prism was used to perform statistical analyses such as unpaired t-test and one-way ANOVA, where appropriate.

---

**Extended Data Fig. 1.**
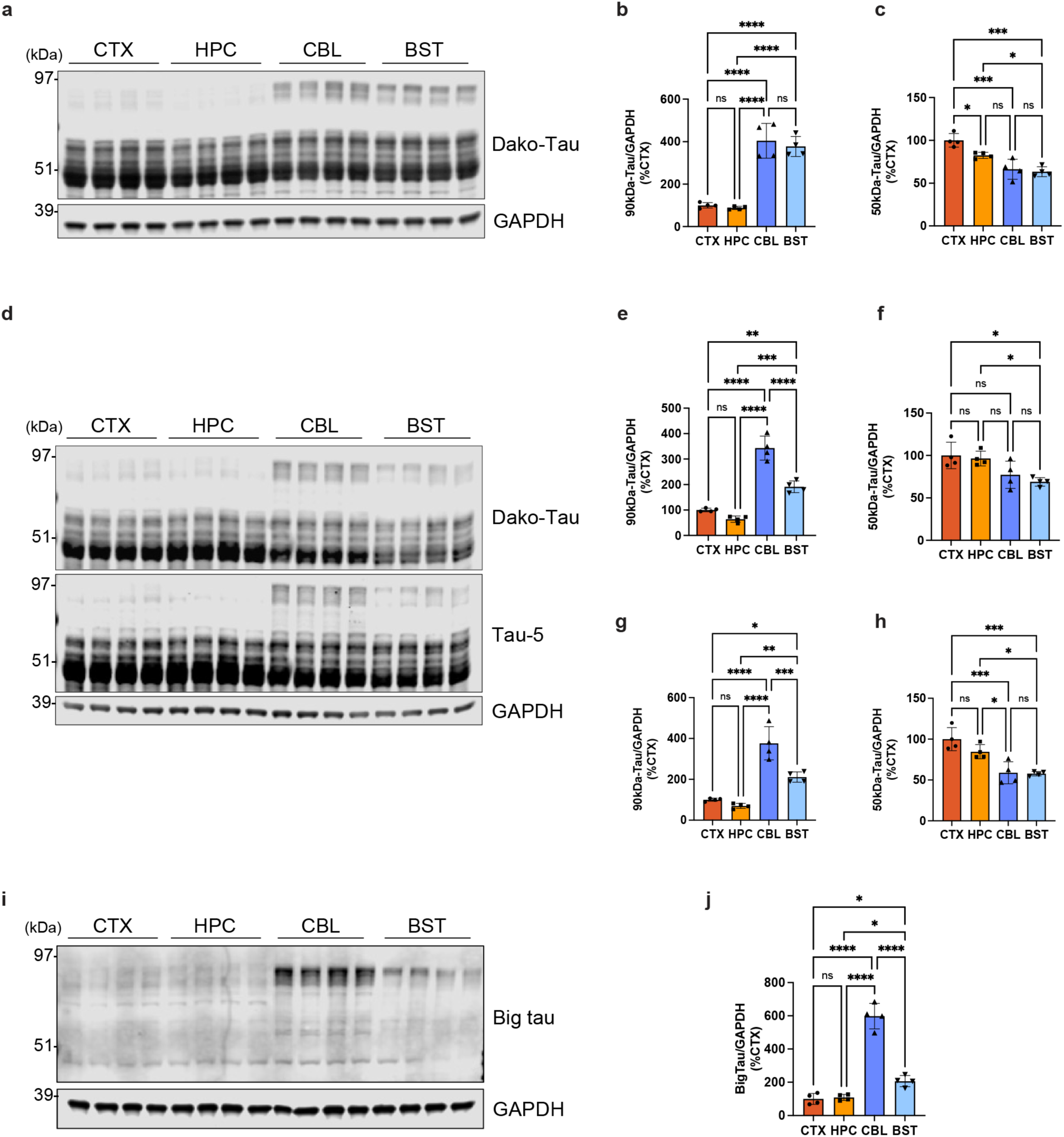
The elevated level of big tau in the cerebellum and brainstem is also detected with a different tau antibody and in the different mouse strain. **a-c,** Representative western blots probed with the Dako-tau antibody showing tau protein expression across different brain regions of 4-6-month-old (mo) C57/B6 mice (**a**). Graphs showing the quantification of tau bands at ∼90kDa (**b**) and ∼50kDa (**c**) normalized to GAPDH. **d-h,** Representative western blots probed with the tau-5 or Dako-tau antibody showing tau protein expression across different brain regions of 1-3-mo FVB mice. (**d**) Graphs showing the quantification of tau bands detected with the tau-5 antibody at ∼90kDa (**e**) and ∼50kDa (**f**), or detected with the Dako-tau antibody at ∼90kDa (**g**) and ∼50kDa (**h**), all normalized to GAPDH. **i-j,** Representative western blots probed with a novel mouse big tau antibody showing the expression of big tau protein in the cerebellum and brainstem of 2-mo FVB mice (**i**) and their quantification normalized to GAPDH (**j**). Data (n=4/group) are shown as mean ± SEM. CTX = cortex; HPC = hippocampus; CBL = cerebellum; BST = brainstem; ns = not significant; \**p*<0.05; \*\**p*<0.01; \*\*\**p*<0.001; \*\*\*\**p*<0.0001; one-way ANOVA.

**Extended Data Fig. 2.**
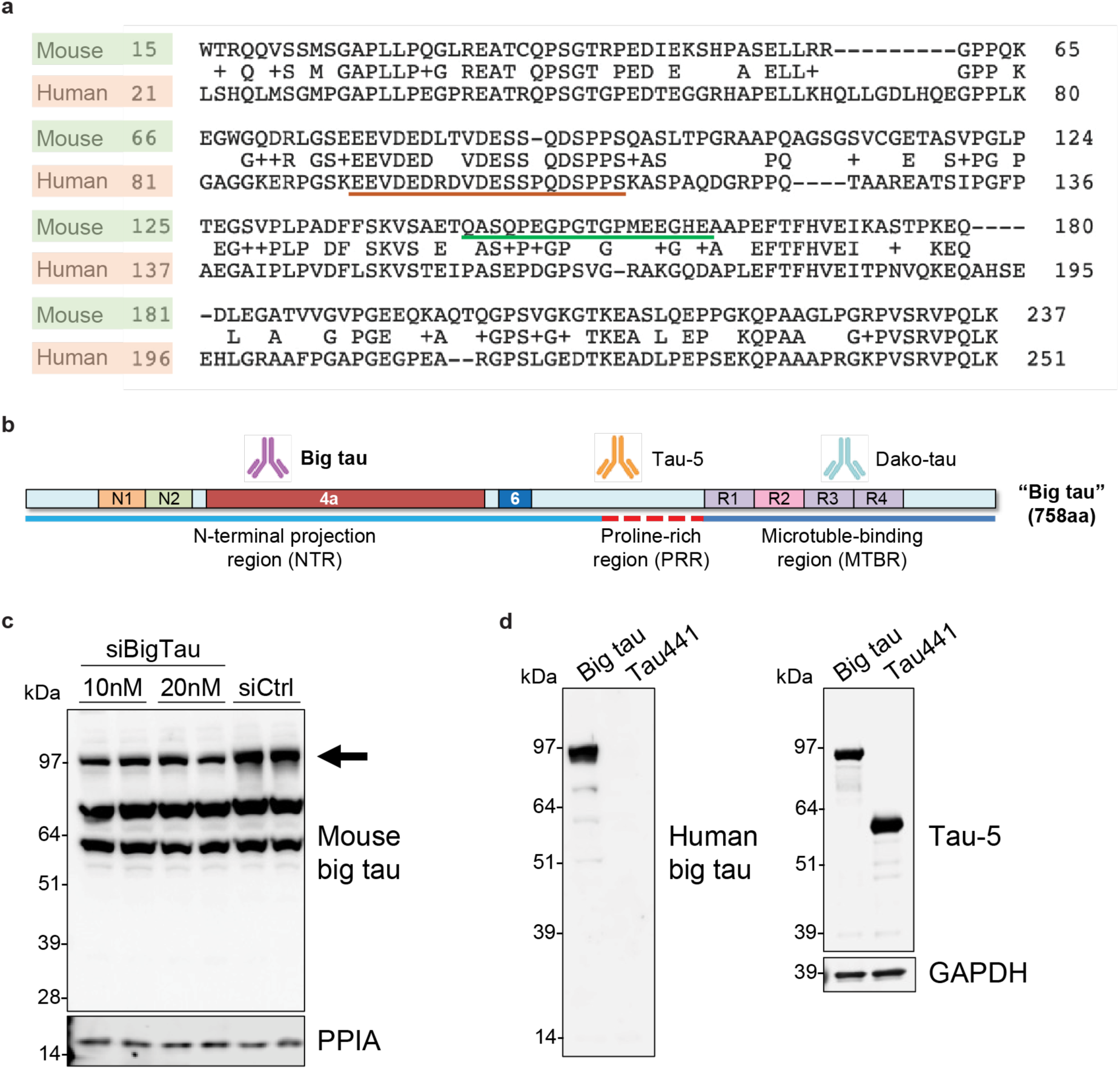
Antibodies that can specifically detect mouse or human big tau protein are generated. **a,** Amino acid sequence comparison between mouse and human big tau proteins. Each peptide (underlined) was selected based on the sequence uniqueness to generate polyclonal antibodies specific to mouse or human big tau protein, respectively. **b,** A schematic of big tau protein (human) depicting the approximate location of the epitope detected by novel big tau antibodies relative to tau-5 and Dako-tau antibodies. **c,** Representative western blots probed with the mouse big tau antibody showing knockdown of big tau (indicated by an arrow) by big tau-targeting siRNA (siBigTau) in the mouse neuroblastoma N1E-115 cell line which expresses endogenous big tau protein^14^. For the negative control, cells were treated with a non-targeting control siRNA (siCtrl) at 10 nM. PPIA was used as a loading control. **d,** Representative western blots probed with the human big tau antibody showing the expression of overexpressed human big tau in HEK293T cells. Tau441 overexpressed in HEK293T cells is detected by the tau-5 antibody but not by the big tau antibody. GAPDH was used as a loading control.

**Extended Data Fig. 3.**
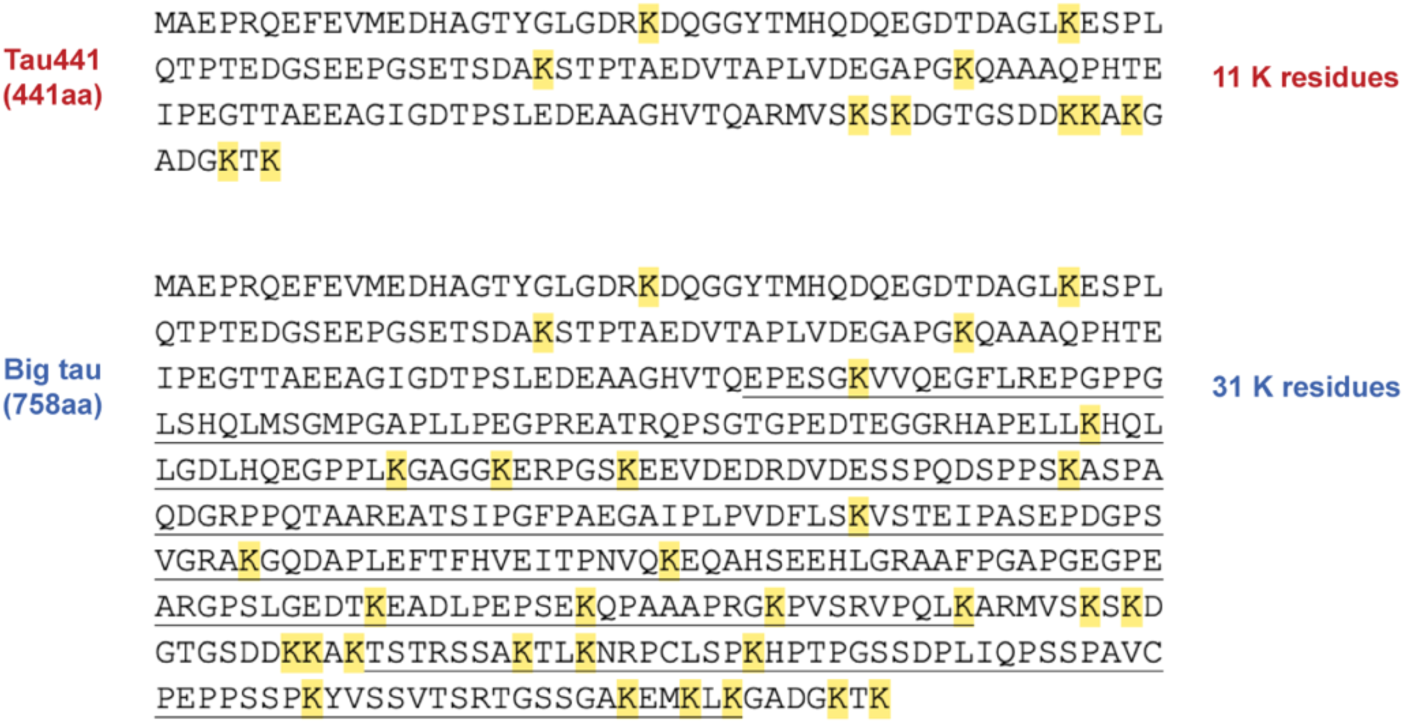
Big tau has 20 more lysine residues than the longest regular tau isoform. Amino acid sequences within the N-terminal projection region (NTR) of tau441 and big tau. Tau441 includes 11 lysine (K) residues whereas big tau includes 31 K residues. K residues are highlighted in yellow.

**Extended Data Fig. 4.**
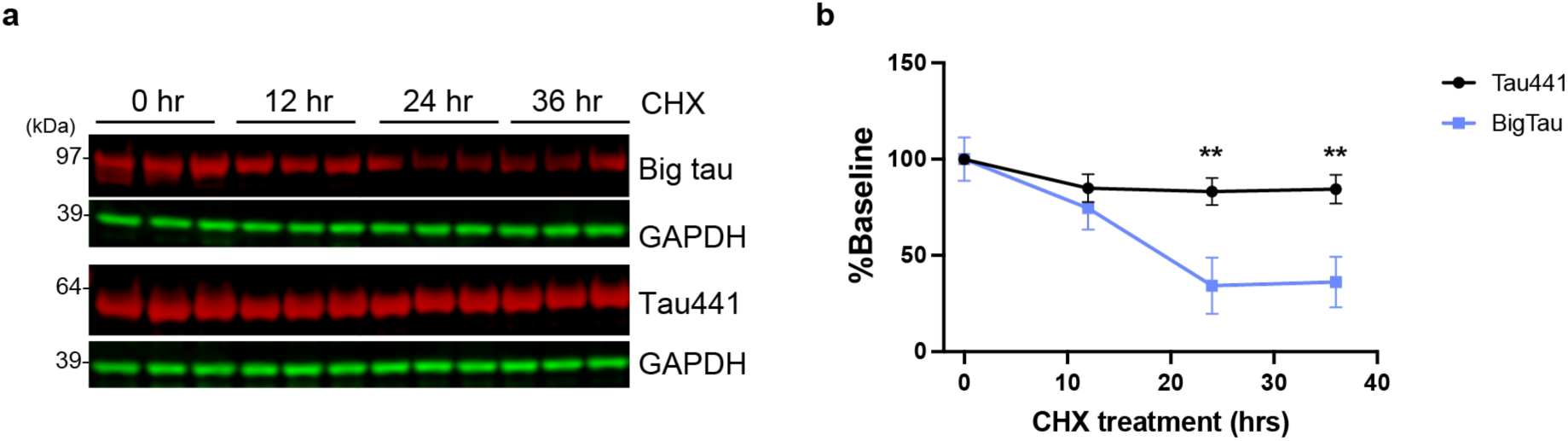
Big tau degrades much more rapidly than tau441. **a-b,** Representative western blots probed with the Dako-tau antibody (**a**) and their quantification normalized to GAPDH (**b**) from the cycloheximide (CHX) chase assay depicting the degree of protein degradation rate between big tau and tau441. N**p<0.01; multiple unpaired t-test.

**Extended Data Fig. 5.**
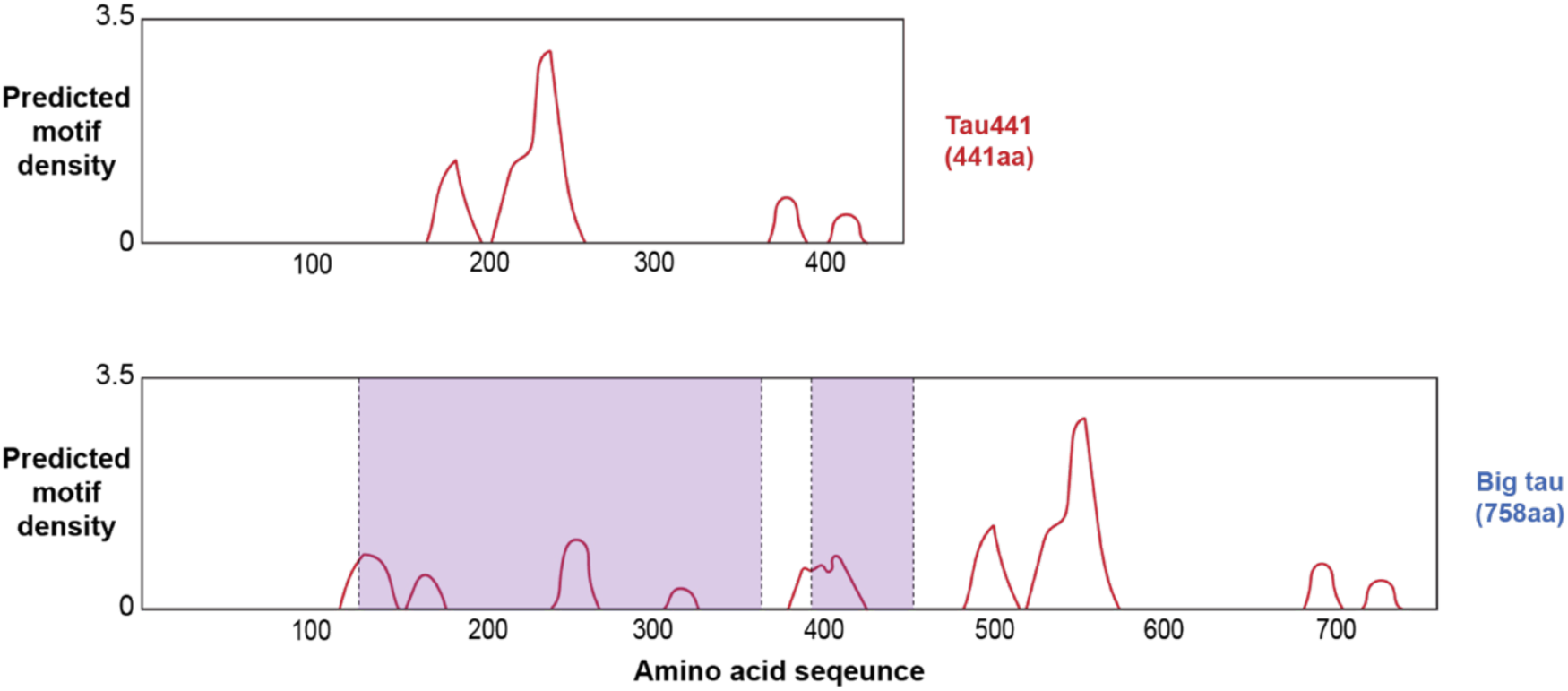
Big tau has additional motifs predicted to bind to microtubules. Graphs showing motifs in tau441 and big tau that are predicted to have binding affinities to microtubules as analyzed by the MAPanalyzer^24^. Purple = regions specific to big tau (absent in regular tau isoforms).

**Extended Data Fig. 6.**
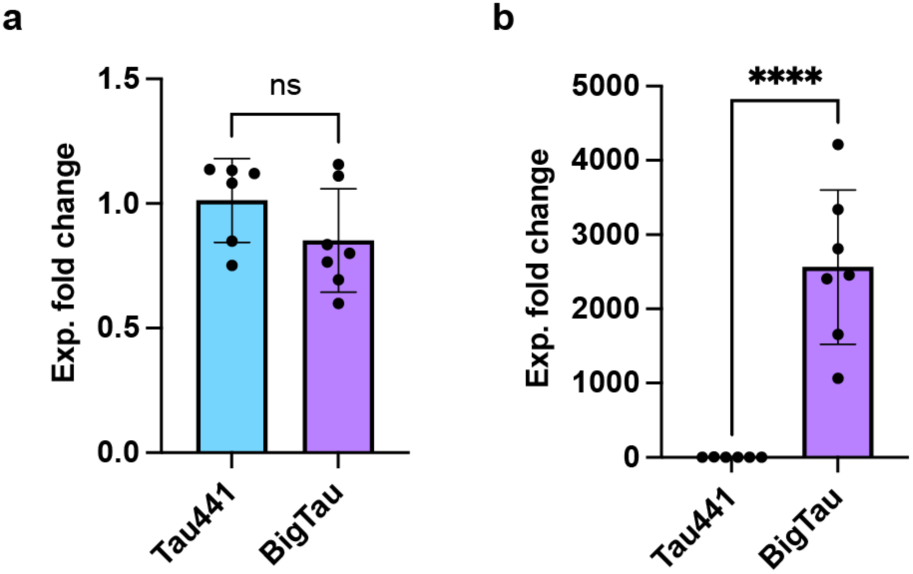
Comparable amounts of tau441 and big tau were expressed in the AAV mouse model. **a-b,** Graphs of qPCR analysis results measuring the mRNA level of total human tau (**a**) or human big tau (**b**) expressed in the forebrain of AAV-injected mice. Data (n=6-7/group) are shown as mean ± SEM. ns = not significant; ****p<0.0001; unpaired t-test.

**Extended Data Fig. 7.**
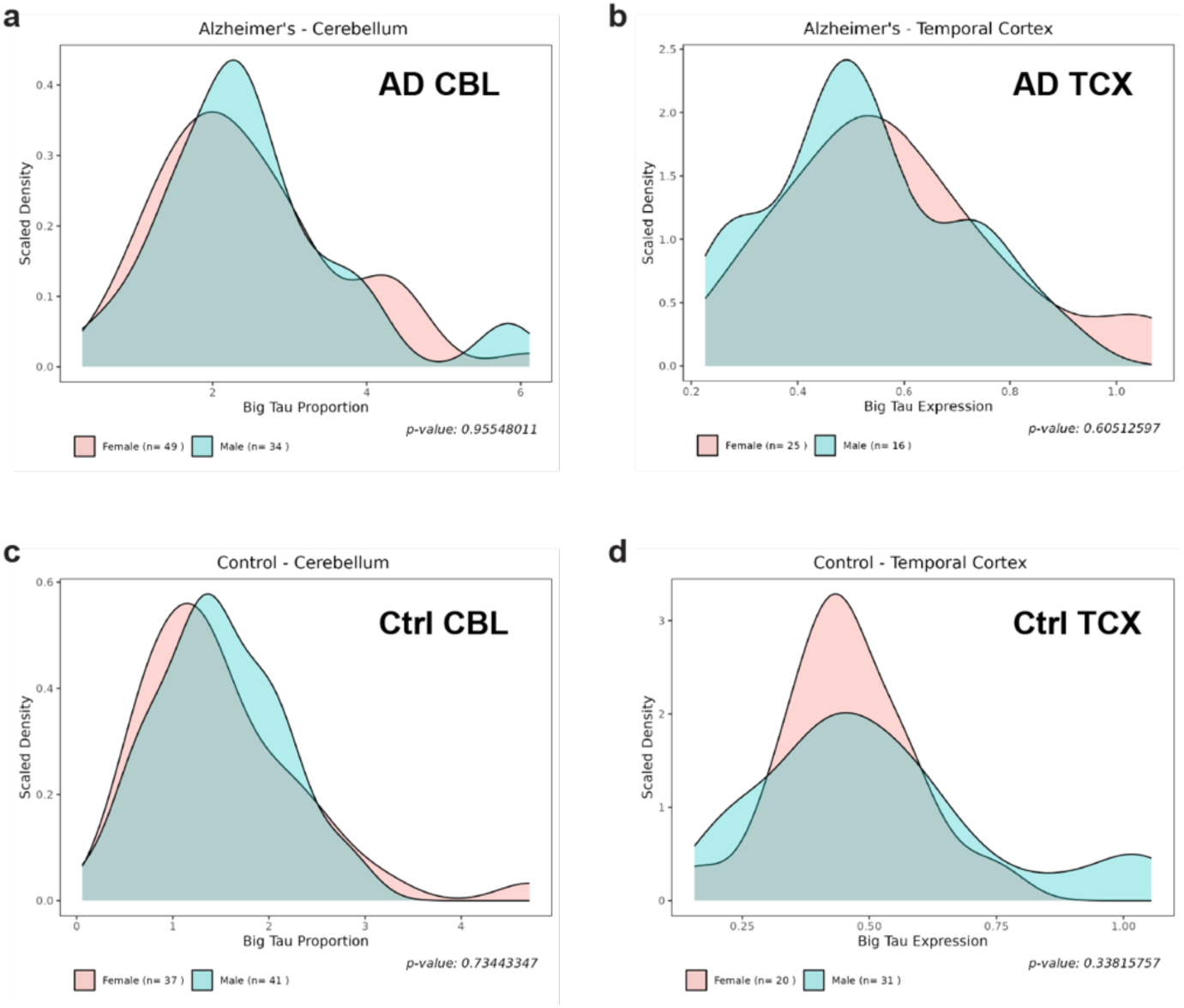
No sex difference is detected in the big tau level per group. **a,** Comparison of the big tau transcript level in the cerebellum among AD patients between females and males. **b,** Comparison of the big tau transcript level in the temporal cortex among AD patients between females and males. **c,** Comparison of the big tau transcript level in the cerebellum among cognitively normal control individuals (ctrl) between females and males. **d,** Comparison of the big tau transcript level in the temporal cortex among cognitively normal control individuals between females and males. CBL = cerebellum. TCX = temporal cortex.

**Extended Data Fig. 8.**
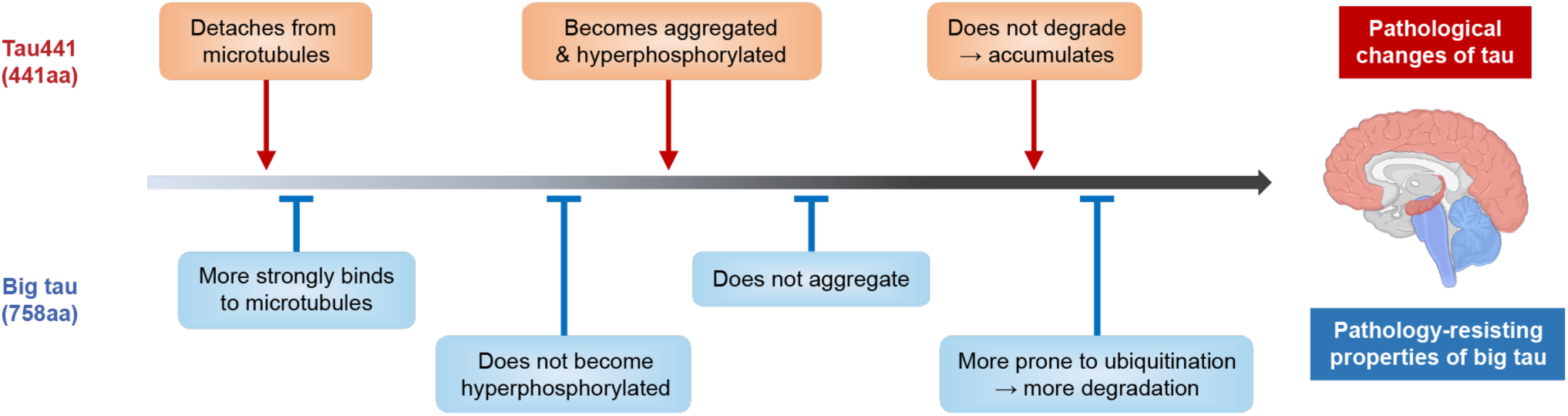
Big tau possesses several pathology-resisting properties. A schematic diagram describing key pathological changes of regular tau isoforms in vulnerable brain regions (shaded in red) during the progression of Alzheimer’s disease (AD) contrasted by pathology-resisting properties of big tau that is highly expressed in brain regions without over tau pathology (shaded in blue). Figures describing each property of big tau are denoted in grey font.

**Extended Data Table 1.** Demographics and pathological information of post-mortem human brain samples used in the study. NPDX = neuropathological diagnosis; PMI = postmortem interval (hours); AD = Alzheimer’s disease case; F = female; M = male; MTG = middle temporal gyrus; CBL = cerebellum.

| Sample | NPDX | Braak | Thal | Age at death (yr) | Sex | PMI | Brain region |
| --- | --- | --- | --- | --- | --- | --- | --- |
| Ctrl 1 | Normal | II | 0 | 79 | F | 9 | CBL |
| Ctrl 2 | Normal | III | 4 | >=90 | F | 18 | MTG, CBL |
| Ctrl 3 | Normal | II | 3 | 89 | M | 36 | MTG, CBL |
| Ctrl 4 | Normal | I | 0 | >=90 | M | 86 | MTG, CBL |
| Ctrl 5 | Normal | I | 0 | 89 | F | 8 | MTG, CBL |
| Ctrl 6 | Normal | 0 | 0 | 59 | M | 35 | CBL |
| Ctrl 7 | Normal | 0 | 0 | >=90 | M | 26 | MTG, CBL |
| Ctrl 8 | Normal | IV | 1 | >=90 | F | 35 | MTG, CBL |
| AD 1 | AD | VI | 5 | 61 | F | 18 | MTG, CBL |
| AD 2 | AD | VI | 5 | 77 | M | 24 | MTG, CBL |
| AD 3 | AD | VI | 5 | 73 | F | 12 | MTG, CBL |
| AD 4 | AD | VI | 5 | 86 | F | 18 | MTG, CBL |
| AD 5 | AD | VI | 5 | 77 | F | 28 | MTG, CBL |
| AD 6 | AD | VI | 5 | 70 | F | 24 | MTG, CBL |
| AD 7 | AD | VI | 5 | >=90 | M | 72 | MTG, CBL |
| AD 8 | AD | VI | 5 | 75 | M | 4 | MTG, CBL |
| AD 9 | AD | VI | 5 | 58 | F | 63 | MTG, CBL |
| AD 10 | AD | VI | 5 | >=90 | M | 6 | MTG, CBL |
| AD 11 | AD | VI | 5 | 75 | F | 26 | MTG, CBL |

